# Lack of selectivity for syntax relative to word meanings throughout the language network

**DOI:** 10.1101/477851

**Authors:** Evelina Fedorenko, Idan Blank, Matthew Siegelman, Zachary Mineroff

**Affiliations:** Department of Brain and Cognitive Sciences, MIT, Cambridge, MA 02139, USA; McGovern Institute for Brain Research, MIT, Cambridge, MA 02139, USA; Department of Psychiatry, MGH, Charlestown, MA 02129, USA; Department of Psychology, UCLA, Los Angeles, CA 90095 USA; Department of Psychology, Columbia University, New York, NY 10027, USA; Eberly Center for Teaching Excellence & Educational Innovation, CMU, Pittsburgh, PA 15213, USA

**Author notes:** Corresponding Author: Ev Fedorenko,; 43 Vassar Street, Room 46-3037G, Cambridge, MA, 02139.

## Abstract

To understand what you are reading now, your mind retrieves the meanings of words and constructions from a linguistic knowledge store (lexico-semantic processing) and identifies the relationships among them to construct a complex meaning (syntactic or combinatorial processing). Do these two sets of processes rely on distinct, specialized mechanisms or, rather, share a common pool of resources? Linguistic theorizing, empirical evidence from language acquisition and processing, and computational modeling have jointly painted a picture whereby lexico-semantic and syntactic processing are deeply inter-connected and perhaps not separable. In contrast, many current proposals of the neural architecture of language continue to endorse a view whereby certain brain regions selectively support syntactic/combinatorial processing, although the locus of such “syntactic hub”, and its nature, vary across proposals. Here, we searched for selectivity for syntactic over lexico-semantic processing using a powerful individual-subjects fMRI approach across three sentence comprehension paradigms that have been used in prior work to argue for such selectivity: responses to lexico-semantic *vs.* morpho-syntactic violations (**Experiment 1**); recovery from neural suppression across pairs of sentences differing in only lexical items *vs.* only syntactic structure (**Experiment 2**); and same/different meaning judgments on such sentence pairs (**Experiment 3**). Across experiments, both lexico-semantic and syntactic conditions elicited robust responses throughout the left fronto-temporal language network. Critically, however, no regions were more strongly engaged by syntactic than lexico-semantic processing, although some regions showed the opposite pattern. Thus, contra many current proposals of the neural architecture of language, syntactic/combinatorial processing is not separable from lexico-semantic processing at the level of brain regions—or even voxel subsets—within the language network, in line with strong integration between these two processes that has been consistently observed in behavioral and computational language research. The results further suggest that the language network may be generally more strongly concerned with meaning than syntactic form, in line with the primary function of language—to share meanings across minds.

## Introduction

### The functional architecture of language processing

What is the functional architecture of human language? A core component is a set of knowledge representations, which include knowledge of words/constructions and their meanings, and the probabilistic constraints on how these linguistic elements can combine to create compound words, phrases, and sentences. During comprehension (decoding of linguistic utterances), we look for matches between elements/substrings in the incoming linguistic signal and these stored knowledge representations, and compose these retrieved representations, in an attempt to re-construct the intended meaning, and during production (encoding of linguistic utterances), we search our knowledge store for the right words/constructions and combine and arrange them in a particular way to express a target idea.

How is this rich set of representations and computations structured? Specifically, which aspects of language are functionally dissociable from one another? Traditionally, two principal distinctions have been drawn: one is between words (the lexicon) and rules (the grammar) (e.g., Chomsky, 1965, 1995; Fodor, 1983; Pinker & Prince, 1988; Pinker, 1991, 1999); and another is between linguistic representations themselves (i.e., our knowledge of the language) and their online processing (i.e., accessing them from memory and combining them to create new complex meanings and structures) (e.g., Chomsky, 1965; Fodor et al., 1974; Newmeyer, 2003). Because these dimensions are, in principle, orthogonal, we could have distinct mental capacities associated with i) knowledge of word (lexical) meanings, ii) knowledge of grammar (syntactic/combinatorial rules), iii) access (or “retrieval”) of lexical representations, iv) access of syntactic/combinatorial rules, and v) combining retrieved representations into new complex representations (**Fig. 1a**).

**Figure 1:**
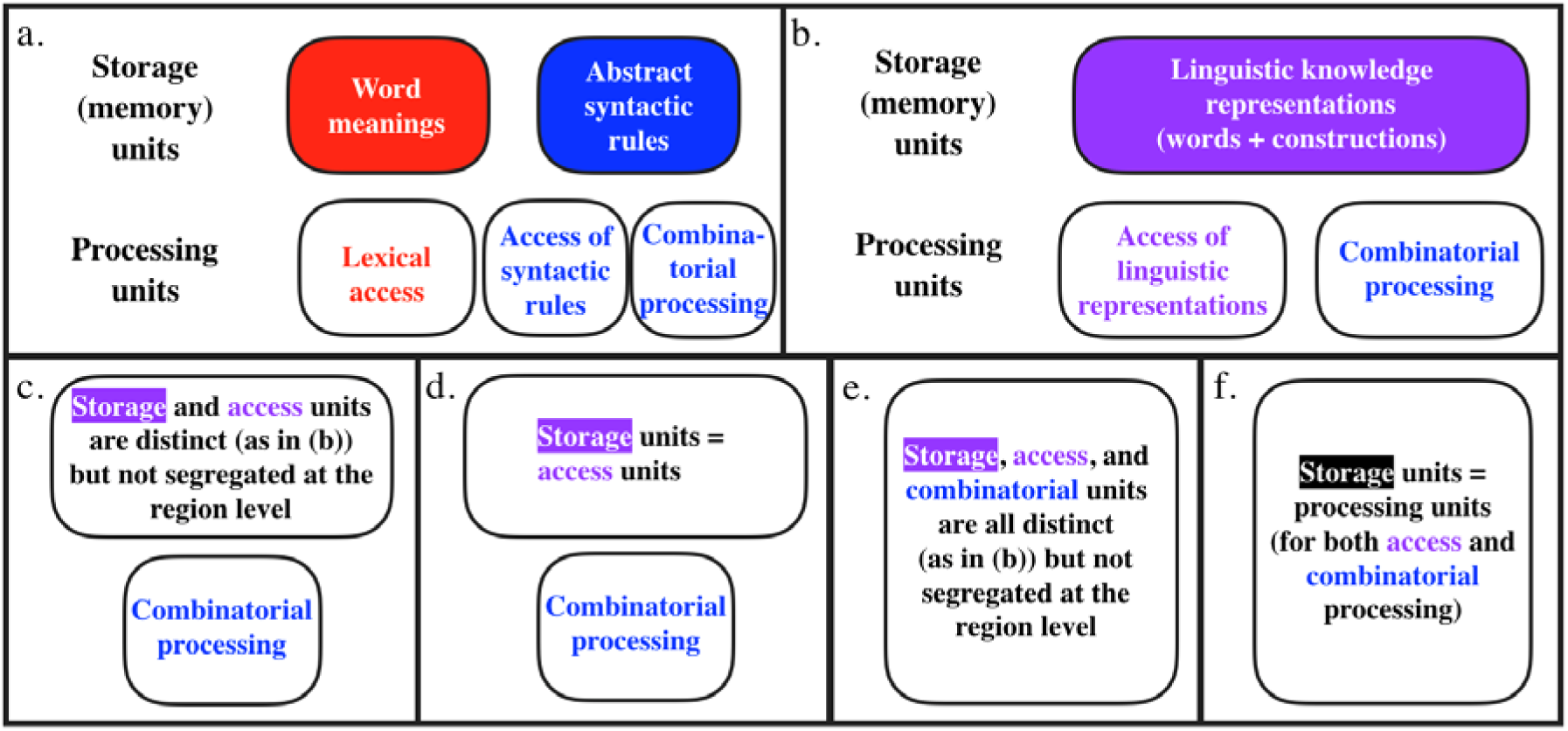
A (non-exhaustive) set of theoretically possible architectures of language. Distinct boxes correspond to distinct brain regions (or sets of brain regions; e.g., in 1a-d, “combinatorial processing” may recruit a single region or multiple regions, but critically, this region or these regions do not support other aspects of language processing, like understanding word meanings). The architectures differ in whether they draw a (region-level) distinction between the lexicon and grammar (a vs. b-f), between storage and access of linguistic representations (1a-b vs. 1c-f), and critically, in whether combinatorial processing is a separable component (1a-d vs. 1e-f).

However, both of these distinctions have been long debated. For example, as linguistic theorizing evolved and experimental evidence accumulated through the 1970s-90s, the distinction between the lexicon and grammar began to blur, both for the storage of linguistic knowledge representations and for online processing (e.g., **Fig. 1b**; see Snider & Arnon, 2012, for a summary and discussion). Many have observed that much of our grammatical knowledge does not operate over general categories like nouns and verbs, but instead requires reference to particular words or word classes (e.g., verbs that can occur in a particular construction, like the ditransitive) (e.g., Lakoff, 1970; Bybee, 1985, 1998, 2010; Levin, 1993; Goldberg, 1995, 2002; Jackendoff, 2002a,b, 2007; Sag et al., 2003; Culicover & Jackendoff, 2005; Levin & Rappaport Hovav, 2005; Audring & Jackendoff, 2020). As a result, current linguistic frameworks incorporate knowledge of “rules” (i.e., syntactic structures) into the mental lexicon, although they differ as to the degree of abstraction that exists above and beyond knowledge of how particular words combine with other words (e.g., Ambridge, 2018; see Hudson, 2007, for discussion), and in whether abstract syntactic representations (like the double object, passive, or question constructions) are always associated with meanings or functions (e.g., Pinker, 1989; Goldberg, 1995; cf. Chomsky, 1957; Branigan & Pickering, 2017; see Jackendoff, 2002, for discussion).

In line with these changes in linguistic theorizing, experimental and corpus work in psycholinguistics have established that humans i) are exquisitely sensitive to contingencies between particular words and the constructions they occur in (e.g., Clifton et al., 1984; MacDonald et al., 1994; Trueswell et al., 1994; Garnsey et al., 1997; Traxler et al., 2002; Reali & Christiansen, 2007; Roland et al., 2007; Jaeger, 2010), and ii) store not just atomic elements (like morphemes and non-compositional lexical items), but also compositional phrases (e.g., “I don’t know” or “give me a break”; e.g., Wray, 2005; Evert, 2008; Arnon & Snyder, 2010; Morgan & Levy, 2016; Christiansen & Arnon, 2017) and constructions (e.g., “the X-er the Y-er”; Goldberg, 1995; Culicover & Jackendoff, 1999). The latter suggested that the linguistic units people store are determined not by their nature (i.e., atomic vs. not) but instead, by their patterns of usage (e.g., Bybee 1998, 2006; Goldberg 2006; Barlow and Kemmer 2000; Langacker 1986, 1987; Tomasello 2003). Further, people’s lexical abilities have been shown to strongly correlate with their grammatical abilities—above and beyond shared variance due to general fluid intelligence—both developmentally (e.g., Bates et al., 1988, 1995; Bates & Goodman, 1997; Dale et al., 2010; Dixon & Marchman, 2007; Snedeker et al., 2007, 2012; Hoff et al., 2018) and in adulthood (e.g., Dabrowska, 2018). Thus, linguistic mechanisms that have been previously proposed to be distinct are instead tightly integrated or, perhaps, are so cognitively inseparable as to be considered a single apparatus.

The distinction between stored knowledge representations and online computations has also been questioned (see Hasson et al., 2015, for a discussion of this issue in language and other domains). For example, by using the same artificial network to represent all linguistic experience, connectionist models dispense not only with the lexicon-grammar distinction but also the storage-computation one, and assume that the very same units that represent our linguistic knowledge support its online access and processing (e.g., Rumelhart & McClelland, 1986; Seidenberg, 1994; Devlin et al., 2019; see also Goldinger, 1996; Bod, 1998, 2006, for exemplar models, which also abandon the storage-computation divide).

### Syntax selectivity in prior cognitive neuroscience investigations?

Alongside psycholinguistic studies, which inform debates about linguistic architecture by examining the behaviors generated by language mechanisms, and computational work, which aims to approximate human linguistic behavior using formal models, a different, complementary approach is offered by cognitive neuroscience studies. These studies aim to constrain the cognitive architecture by examining how cognitive processes are neurally implemented (e.g., Kanwisher, 2010; Mather et al., 2013). The assumption that links neuroimaging data (and neuropsychological patient data) to cognitive hypotheses is as follows: if distinct brain regions or sets of regions support the processing of manipulations targeting cognitive processes X and Y, we can infer that X and Y are dissociable. Such brain regions would be expected to show distinct patterns of response in brain imaging studies, and their damage should lead to distinct patterns of cognitive deficits.

A large number of brain imaging investigations have observed distinct loci of activation for manipulations that target (lexico-)semantic vs. syntactic processing, and—following the reasoning above—have argued for a dissociation between the two (e.g., Dapretto & Bookheimer, 1999; Embick et al., 2000; Kuperberg et al., 2000; Ni et al., 2000; Newman et al., 2001; Kuperberg et al., 2003; Noppeney & Price, 2004; Cooke et al., 2006; Friederici et al., 2010; Glaser et al., 2013; Schell et al., 2017, inter alia). Consequently, many proposals of the neural architecture of language postulate a component that selectively supports syntactic, or more general combinatorial, processing relative to the storage/processing of individual word meanings (e.g., Grodzinsky & Santi, 2008; Baggio and Hagoort, 2011; Friederici, 2011, 2012; Tyler et al., 2011; Duffau et al., 2014; Ullman, 2016; Matchin & Hickok, 2019; Pylkkanen, 2019; cf. Bornkessel-Schlesewsky & Schlesewsky, 2009; Bornkessel-Schlesewsky et al., 2015 – we come back to these proposals in the Discussion). Although proposals vary in which component(s) of syntactic processing are emphasized—from morpho-syntactic agreement, to dependency structure building and composition, to word-order-related processes—the general idea of syntax selectivity remains prominent in cognitive neuroscience of language. How can we fit such selectivity with current linguistic theorizing, psycholinguistic evidence, and computational modeling work, which suggest strong integration between lexico-semantic and syntactic representations and processing, as discussed above? Here, we revisit past evidence and report a new study to ***argue against the existence of brain regions that are selective for syntactic/combinatorial processing over the processing of word meanings***.

There already exist ***reasons to doubt*** the existence of syntax-selective brain regions if we take a closer look at the cognitive neuroscience of language literature. ***First***, the specific brain regions that have been argued to support syntactic/combinatorial processing, and the construal of these regions’ contributions, *differ across studies and theoretical proposals* (e.g., Baggio and Hagoort, 2011; Bemis & Pylkkanen, 2011; Friederici, 2011, 2012; Tyler et al., 2011; Duffau et al., 2014; Ullman, 2004, 2016; Matchin & Hickok, 2019; Pylkkanen, 2019). For example, the proposed location of the “core” syntactic/combinatorial hub varies between the inferior frontal cortex (e.g., Friederici et al., 2006; Hagoort, 2005, 2013), the posterior temporal cortex (e.g., Matchin & Hickok, 2019), and the anterior temporal cortex (e.g., Vandenberghe et al., 2002; Bemis & Pylkkanen, 2011; Pylkannen, 2019), with some positing multiple hubs (e.g., Pallier et al., 2011; Tyler et al., 2011).

***Second***, at least some syntactic/combinatorial manipulations appear to engage the *entire fronto-temporal language network* (e.g., Fedorenko et al., 2010; Pallier et al., 2011; Blank et al., 2016), putting into question the idea of syntactic processing being focally carried out. Relatedly, studies of patients with brain damage have failed to consistently link syntactic deficits with a particular region within the language network. Instead, damage to any component of the network appears to lead to similar syntactic difficulties—which also mirror patterns observed in neurotypical individuals under cognitive load (Miyake et al., 1994)—leading some to argue that syntactic processing is supported by the language network as a whole (e.g., Caplan et al., 1996; Dick et al., 2001; Wilson and Saygin, 2004; Mesulam et al., 2014, 2015). Given that the language network has to additionally support lexico-semantic processing, these findings necessarily imply that at least some, and possibly all, of these syntax-responsive regions overlap with regions that support lexico-semantic processing.

***Third***, a number of studies have actually *failed to observe syntax selectivity*, showing that brain regions that respond to syntactic manipulations also show reliable, and sometimes stronger, responses to lexico-semantic manipulations (e.g., Chee et al., 1999; Keller et al., 2001; Roder et al., 2002; Luke et al., 2002; Heim et al., 2008; Rogalsky & Hickok, 2009; Fedorenko et al., 2010, 2012a, 2016; Bautista & Wilson, 2016; Blank et al., 2016; Shain et al., 2019; see Rodd et al., 2015, for a meta-analysis). Relatedly, some studies have reported activations for lexico-semantic manipulations, such as lexical ambiguity manipulations, in what appear to be the same regions as the ones implicated in other studies in syntactic/combinatorial processing: i.e., regions in the left inferior fontal and posterior temporal cortex (e.g., Rodd et al., 2005, 2010, 2012; Davis et al., 2007; Mason and Just, 2007; Zempleni et al., 2007; Bilenko et al., 2008; Bekinschtein et al., 2011).

And ***fourth***, many prior studies that have reported syntax selectivity suffer from *methodological and statistical limitations*. For example, although diverse paradigms have been used across studies to probe lexico-semantic vs. syntactic/combinatorial processing, any given study has typically used a single paradigm, raising the possibility that the results reflect paradigm-specific differences between conditions rather than a general difference between lexico-semantic and syntactic/combinatorial representations or computations. Furthermore, many studies that claimed to have observed a dissociation have not reported the required region-by-condition interactions, as needed to argue for a functional dissociation between brain regions (Nieuwenhuis et al., 2011). And some studies have argued for syntax selectivity based solely on sensitivity to syntactic complexity manipulations, without even examining responses to lexico-semantic processing (e.g., Stromswold et al., 1996; Ben-Shachar et al., 2003; Fiebach et al., 2005; Santi & Grodzinsky, 2010; see Friederici, 2011, for a meta-analysis). Although such studies (may) establish that a brain region is engaged in syntactic processing, they say little about its *selectivity* for syntactic over lexico-semantic processing. Finally, some studies have reported sensitivity to syntactic manipulations in regions that fall outside the boundaries of the fronto-temporal language network. For example, some studies of morpho-syntactic violations (e.g., Kuperberg et al., 2003; Nieuwland et al., 2012) have reported effects in regions that resemble the domain-general bilateral fronto-parietal network implicated in executive control (e.g., Duncan, 2010, 2013). This network is sensitive to unexpected events across domains (e.g., Corbetta & Shulman, 2002), and some of its regions lie in close proximity to the language regions (e.g., Fedorenko et al., 2012b; Fedorenko & Blank, in press). Although the precise nature of this network’s contribution to language processing remains debated (e.g., Fedorenko, 2014; Diachek et al., 2019; Ryskin et al., 2020a), sensitivity of this network to a linguistic manipulation likely indexes a domain-general process and does not inform the question of whether different components of the *language network* support syntactic vs. lexico-semantic processing.

### Motivation for the current study

The current study aims to ***resolve the conflict*** between i) converging evidence from linguistic theorizing, behavioral psycholinguistic work, and computational modeling, which have jointly painted a clear picture of strong integration between lexico-semantic and syntactic representations and processing, and ii) cognitive neuroscience studies and proposals, many of which continue to suggest the existence of syntax- or combinatorics-selective brain regions. To do so, we use fMRI to search for syntax selectivity using a *robust individual-subjects analytic approach*, including a well-validated task for identifying the language network (i.e., a “functional localizer”; Fedorenko et al., 2010) across *three classic paradigms* from the literature that contrast lexico-semantic and syntactic processing. We will now discuss and motivate both of these features of our study in greater detail, in the historical context of the field.

#### The need for robust, replicable, and cumulative science

Over the years, a large number of paradigms have been used across brain imaging studies to probe lexico-semantic and syntactic processes and their relationship (see references in Materials and Methods). Some paradigms (discussed below) have varied *the presence or absence* of lexico-semantic vs. syntactic information in the linguistic signal; others have *more strongly taxed* the processing of word meanings vs. syntactic structure; and still others have made the meaning of a particular word vs. the structure of the sentence *more salient / task-relevant.* Any of these kinds of manipulations could be informative, but no single manipulation—and certainly not from a single experiment—would suffice to argue for syntax selectivity. To compellingly argue that a brain region selectively supports (some aspect of) syntactic processing, one would need to demonstrate the *robustness* of the syntactic > lexico-semantic effect (e.g., replication in a new sample of participants, on a new set of experimental materials, and/or in another imaging modality) and its *generalizability* to other contrasts between conditions that engage the hypothesized computation and ones that do not. In particular, the selectivity of a brain region for the critical syntactic condition(s) would need to be established relative to a *broad range of control conditions*, given that any given syntactic vs. lexico-semantic pair of conditions will likely differ in ways beyond the target distinction between syntax and semantics. For example, showing that morpho-syntactic violations elicit a stronger response than semantic violations is not sufficient to argue for syntactic selectivity because these two conditions also differ in whether the error can be explained by a plausible noise process within a noisy-channel framework of sentence comprehension (e.g., Ferreira et al., 2002; Levy et al., 2009; Gibson et al., 2013); thus, a stronger response to morpho-syntactic violations could reflect the relevant correction process, which does not get engaged for typical semantic violations (cf. Ryskin et al., 2020b). Furthermore, it would be critical to ensure that—across studies—the critical effect arises in the *same* brain region, not just within the same broad macroanatomic area, like the left inferior frontal gyrus (LIFG) (see Hong et al., 2019, for a general discussion of challenges to determining what counts as a “replication” in fMRI research). To the best of our knowledge, nobody has demonstrated syntactic selectivity using this kind of a rigorous approach, or even attempted to do so. At least two factors have likely contributed to the lack of such attempts, and to the resulting lack of clarity in the field.

First, until a decade ago, the issue of replicability has not been much discussed in the fields of psychology / cognitive science (e.g., Ioannidis et al., 2014) and cognitive neuroscience (e.g., Poldrack et al., 2017). And at least some of the findings in cognitive neuroscience of language that have been taken at face value based on a single report by a single group may not be robust and replicable (e.g., see Siegelman et al., 2019, for a recent attempt, and failure, to replicate a much cited report by Dapretto & Bookheimer, 1999). This issue is further compounded by the many hidden degrees of freedom (e.g., Simmons et al., 2011) that characterize the choices during the preprocessing and analysis of brain imaging data (Botvinik-Nazer et al., 2019) and the common use of “double dipping” (e.g., Krigeskorte et al., 2009) in many early studies. And second, as we have previously argued (e.g., Fedorenko & Kanwisher, 2009; Fedorenko et al., 2010, 2012b; Blank et al., 2017; Fedorenko & Blank, 2020; see also Brett et al., 2002 and Saxe et al., 2006), establishing a cumulative research enterprise in cognitive neuroscience of language has been challenging due to the difficulty of comparing findings across studies that rely on the traditional group-averaging approach (e.g., Holmes & Friston, 1998). In this analytic approach—which has dominated the brain-imaging language research in the 1990s and 2000s and is still in common use despite being strongly disfavored in neuroimaging studies in other domains—individual activation maps are aligned in a common space, and the output is a set of coordinates in that space for voxels where significant effects obtain (typically the most reliable peak(s) in each activation cluster are reported). The main way to compare results across such studies is to compare the anatomical locations of these activation peaks. However, group-level activation peaks are noisy (e.g., Fedorenko et al., 2012b), due to the combined effect of two factors, both especially pronounced in the association cortex, which houses the language system: (1) high inter-individual variability in the locations of functional areas (e.g., Fedorenko et al., 2010; Mahowald & Fedorenko, 2016; Braga et al., 2019); and (2) lack of correspondence between functional areas and macroanatomic landmarks, like gyri and sulci (e.g., Frost & Goebel, 2011; Tahmasebi et al., 2011; Vázquez-Rodríguez et al., 2019). And even the most systematic comparisons in the form of meta-analyses of activation peaks from large numbers of studies (e.g., Bookheimer, 2002; Costafreda et al., 2006; Indefrey & Levelt, 2004; Kaan & Swaab, 2002; Lindenberg et al., 2007; Poldrack et al., 1999; Vigneau et al., 2006; Binder et al., 2009) are not very informative (Kvarven et al., 2019), and have been shown to lead to fundamentally wrong conclusions about the functional architecture of the human brain in some cases (e.g., Aguirre & Farah, 1998). So what is a solution?

A decade ago, we developed an alternative approach to the study of language in the brain (Fedorenko et al., 2010)—one that had been successful in other domains, including high-level vision (e.g., Kanwisher et al., 1997) and social cognition (e.g., Saxe & Kanwisher, 2003), and has now become widespread in the study of language (e.g., Axelrod et al., 2015; Lane et al., 2015; Poldrack et al., 2015; Wang et al., 2015; Matchin et al., 2019; Braga et al., 2019). In this approach, language-responsive areas are defined functionally in individual brains without being constrained to fall precisely in the same anatomical locations across participants; and the localized regions are then probed for their responses to critical experimental manipulations. The use of the same “localizer” paradigm across individuals, studies, and research groups (and in domains, like vision, this is done across species, too; e.g., Tsao et al., 2008) provides a straightforward way to *directly relate findings* to one another.

#### Choice of paradigms

Using the individual-subjects functional localization approach, we have previously argued for the lack of syntax selectivity based on a paradigm that varies the presence of lexico-semantic vs. syntactic information in the linguistic signal (Fedorenko et al., 2010, 2012a, 2016; for earlier uses of this paradigm, see e.g., Mazoyer et al., 1993; Friederici et al., 2000; Humphries et al., 2001; Vandenberghe et al., 2002; for another variant, see Bautista & Wilson, 2016). In particular, we examined the processing of i) sentences, which have a syntactic structure and consist of real, interpretable words, ii) lists of unconnected words, which lack structure but are individually interpretable, iii) “Jabberwocky” sentences, which preserve a syntactic frame (word order and morpho-syntactic endings), but have the words replaced by nonwords, so the meanings of those strings cannot be interpreted with respect to our world knowledge, aside from very coarse-level semantics, and finally, iv) lists of unconnected nonwords, which lack both structure and interpretability. Across three replications with fMRI (Fedorenko et al., 2010; see Mollica et al., in prep. for another replication) and, in addition, in a more spatially and temporally sensitive method (electrocorticography, ECoG) (Fedorenko et al., 2016), we found that any language-responsive brain region or electrode that shows sensitivity to syntactic structure (i.e., stronger responses to sentences than word lists, and to Jabberwocky sentences than nonword lists) is at least as sensitive, and often more sensitive, to meanings of individual words (showing stronger responses to sentences than Jabberwocky sentences, and to word lists than nonword lists).

However, one could question the findings from this paradigm because the contrasts are rather crude and the materials are artificial/unnatural. If the overlap in the brain mechanisms that process individual word meanings and syntactic structure is a real and robust finding, the results should generalize to other, finer-grained comparisons between lexico-semantic and syntactic processing. As a result, we selected three paradigms from studies that have argued for syntax selectivity or for dissociations between lexico-semantic and syntactic processing, and that continue to be cited as evidence of such, and attempted to conceptually replicate (Schmidt, 2009) them.

Experiments 1 and 3 are designed to differentially tax lexico-semantic vs. syntactic processing by having a critical word in a sentence be incompatible with the context in terms of either its its meaning or morpho-syntactic properties (Experiment 1), or by forcing participants to focus on the meanings of the critical words or the structure of sentences (Experiment 3). Experiment 2 relies on the well-established neural adaptation to the repetition of a stimulus and recovery from such adaptation when some relevant feature of the stimulus changes: here, a change in the individual words (but not the sentence structure) vs. the sentence structure (but not the words). Along with the manipulations varying the presence/absence of lexico-semantic and syntactic information in the linguistic signal discussed above, these manipulations span the space of available manipulations targeting lexico-semantic and syntactic processing quite comprehensively1, and—for syntactic processing—cover both morpho-syntactic agreement (Experiment 1), and dependency structure building / word-order-related processes (Experiments 2 and 3).

If *any brain region* within the language network selectively supports syntactic processing, we would expect stronger responses to the syntactic than the lexico-semantic condition in that region in at least one paradigm. If this pattern holds—for the same brain region(s)—across two or all three paradigms, that would further help rule out paradigm-specific between-condition differences/confounds and strengthen the conclusion. Note that unlike the paradigms that vary the presence/absence of syntactic and lexical information in the linguistic signal—where one could, in principle, observe a pattern where a brain region is not *at all* engaged unless structure or meaning is present (although in practice, even lists of pseudowords, which lack both structure and meaning, elicit an above-baseline response across much of the language network; Fedorenko et al., 2010)—the current paradigms all use sentence materials across conditions, so all conditions are expected to elicit above-baseline responses throughout the language network. The critical question is whether any brain region(s) would exhibit stronger responses for the syntactic compared to lexico-semantic condition in one or more paradigms. If no brain region within the language network shows this pattern, this would strongly reinforce the conclusions drawn from the paradigms that have varied the presence/absence of lexico-semantic and syntactic information in the signal (e.g., Fedorenko et al., 2010; Bautista & Wilson, 2016).

To foreshadow the results: using the most sensitive analytic methods available in fMRI (e.g., Nieto-Castañon & Fedorenko, 2012), we find robust responses to both lexico-semantic and syntactic processing throughout the language network in each of the three experiments. Critically, every brain region in the language network that responds to syntactic manipulations responds at least as strongly to lexico-semantic manipulations. No region—or even set of non-contiguous voxels within these regions—shows a consistent preference, in the form of a stronger response, for syntactic processing (ruling out architectures in **Figure 1a-d**). However, in line with our prior work (e.g., Fedorenko et al., 2012a, 2016), some regions show the opposite preference—a stronger response to lexico-semantic processing. We therefore hope that this study brings clarity to the field and helps build stronger bridges, grounded in robust empirical work, with behavioral and computational investigations of language processing.

## Materials and Methods

### General description of the paradigms and their use in prior studies

The *first paradigm* is commonly used in ERP investigations of language processing and relies on violations of expectations about an incoming word that are set up by the preceding context. In particular, the critical word does not conform to either the lexico-semantic or the syntactic expectations (e.g., Kutas & Hilliyard, 1980; Osterhout & Holcomb, 1992; Hagoort et al., 1993). This paradigm has been used in a number of prior fMRI studies (e.g., Embick et al., 2000; Newman et al., 2001; Kuperberg et al., 2003; Cooke et al., 2006; Friederici et al., 2010; Herrmann et al., 2012). The *second paradigm* relies on neural adaptation, wherein repeated exposure to a stimulus leads to a reduction in response, and a change in some feature(s) of the stimulus leads to a recovery of response (see e.g., Krekelberg et al., 2006, for a general overview of the approach). This paradigm has also been used in prior fMRI studies that examined adaptation, or recovery from adaptation, to the lexico-semantic vs. syntactic features of linguistic stimuli (e.g., Noppeney & Price, 2004; Devauchelle et al., 2009; Santi & Grodzinsky, 2010; Menenti et al., 2012; Segaert et al., 2012). Finally, the *third paradigm* was introduced in a classic study by Dapretto & Bookheimer (1999): pairs of sentences vary in a single word vs. in word order / syntactic structure. Either of these manipulations can result in the same meaning being expressed across sentences (if a word is replaced by a synonym, or if a syntactic alternation, like active➔passive, is used) or in a different meaning (if a word is replaced by a non-synonym, or if the thematic roles are reversed). Participants make same/different meaning judgments on the resulting sentence pairs.

### General analytic approaches

In an effort to maximize sensitivity and functional resolution (e.g., Nieto-Castañon & Fedorenko, 2012), we adopt an approach where all the key contrasts are performed within individual participants. We perform two kinds of analyses. In one set of analyses, we identify language-responsive areas in each individual participant with an independent language “localizer” task based on a contrast between the reading of sentences vs. sequences of nonwords (Fedorenko et al., 2010), and compare the responses of these areas to the lexico-semantic vs. syntactic conditions in each critical paradigm. As discussed in the Introduction, the use of the same language localizer task across experiments allows for a straightforward comparison of their results, obviating the need to rely on coarse anatomy and reverse-inference reasoning (Poldrack, 2006, 2011) for interpreting activations in functional terms.

To further ensure that we are not missing syntax selectivity by averaging across (relatively) large sets of language-responsive voxels (see Friston et al., 2006, for discussion of this potential issue), we supplement these analyses with analyses where we only use data from the critical paradigms. In particular, we use some of the data from a given critical task to search for the most syntactic-selective (or lexico-semantic-selective, examined for completeness) voxels (e.g., in Experiment 1, voxels that respond more strongly to morpho-syntactic than semantic violations), and then test the replicability of this selectivity in left-out data, as detailed below.

### Participants

Forty-nine individuals (age 19-32, 27 females) participated for payment (Experiment 1: n=22; Experiment 2: n=14; and Experiment 3: n=15; 2 participants overlapped between Experiments 1 and 3). Forty-seven were right-handed, as determined by the Edinburgh handedness inventory (Oldfield, 1971), or self-report; the two left-handed individuals showed typical left-lateralized language activations in the language localizer task (see Willems et al., 2014, for arguments to include left-handers in cognitive neuroscience research). All participants were native speakers of English from Cambridge, MA and the surrounding community. One additional participant was scanned (for Experiment 2) but excluded from the analyses due to excessive sleepiness and poor behavioral performance. All participants gave informed consent in accordance with the requirements of MIT’s Committee on the Use of Humans as Experimental Subjects (COUHES).

### Design, stimuli, and procedure

Each participant completed a language localizer task (Fedorenko et al., 2010) and a critical task (35 participants performed the localizer task in the same session as the critical task, the remaining 14 performed the localizer in an earlier session; see Mahowald & Fedorenko, 2016 and Braga et al., 2019, for evidence that localizer activations are stable across scanning sessions). Most participants completed one or two additional tasks for unrelated studies. The entire scanning session lasted approximately 2 hours.

#### Language localizer task

The task used to localize the language network is described in detail in Fedorenko et al. (2010). Briefly, we used a reading task that contrasted sentences and lists of unconnected, pronounceable nonwords in a standard blocked design with a counterbalanced order across runs (for timing parameters, see **Table 1**). This contrast targets higher-level aspects of language including, critically, both lexico-semantic and syntactic/combinatorial processing, to the exclusion of perceptual (speech or reading-related) and articulatory processes (see e.g., Fedorenko & Thompson-Schill, 2014, for discussion). As discussed above, brain regions thus localized indeed show sensitivity to lexico-semantic processing (as evidenced by stronger responses to sentences than Jabberwocky sentences, and to word lists than to nonword lists) and to syntactic processing (as evidenced by stronger responses to sentences that word lists, and to Jabberwocky sentences than nonword lists) (e.g., Fedorenko et al., 2010, 2012a, 2016; Mollica et al., in prep.). Stimuli were presented one word/nonword at a time. For 19 participants, each trial ended with a memory probe and they had to indicate, via a button press, whether or not that probe had appeared in the preceding sequence of words/nonwords. The remaining 30 participants read the materials passively and performed a simple button-press task at the end of each trial (included in order to help participants remain alert). Importantly, this localizer has been shown to generalize across different versions: the sentences > nonwords contrast, and similar contrasts between language and a degraded control condition, robustly activates the fronto-temporal language network regardless of the task, materials, and modality of presentation (Fedorenko et al., 2010; Fedorenko, 2014; Scott et al., 2016). Furthermore, the same network robustly emerges from naturalistic-cognition paradigms (e.g., resting state, listening to stories, watching movies) using the functional correlation approach (e.g., Blank et al., 2014; Tie et al., 2014; Branco et al., 2019;Braga et al., 2019), suggesting that this network constitutes a natural kind in the brain, and our localizer contrast is simply a quick and efficient way to identify the relevant areas.

**Table 1.** Timing parameters for the different versions of the language localizer task.

|  | Version |  |  |
| --- | --- | --- | --- |
|  | A | B | C |
| Number of participants | 30 | 14 | 5 |
| Task: Button Press or Memory? | BP | M | M |
| Words / nonwords per trial | 12 | 12 | 12 |
| Trial duration (ms) | 6,000 | 6,000 | 6,000 |
| Fixation | 100 | --- | --- |
| Presentation of each word / nonword | 450 | 350 | 350 |
| Fixation | --- | 300 | 300 |
| Button icon / Memory probe | 400 | 1,000 | 1,000 |
| Fixation | 100 | 500 | 500 |
| Trials per block | 3 | 3 | 3 |
| Block duration (s) | 18 | 18 | 18 |
| Blocks per condition (per run) | 8 | 8 | 6 |
| Conditions | Sentences | Sentences | Sentences |
|  | Nonwords | Nonwords | Nonwords |
|  |  |  | Word-lists* |
| Fixation block duration (s) | 14 | 18 | 18 |
| Number of fixation blocks | 5 | 5 | 4 |
| Total run time (s) | 358 | 378 | 396 |
| Number of runs | 2 | 2 | 2-3 |

\* Used for an unrelated experiment.

#### Critical experiments

The key details about the three experiments are presented in **Figure 2** (sample stimuli), **Figure 3** (trial structure), and **Table 2** (partitioning of stimuli into experimental lists and runs). To construct experimental stimuli, we first generated for each experiment a set of “base items” and then edited each base item to create several, distinct versions corresponding to different experimental conditions. The resulting stimuli were divided into several experimental lists following a Latin Square design, such that in each list (i) stimuli were evenly split across experimental conditions, and (ii) only one version of each item was used. Each participant saw materials from a single list, divided into a few experimental runs. All experiments used an event-related design. Condition orders were determined with the optseq2 algorithm (Dale, 1999), which was also used to distribute inter-trial fixation periods so as to optimize our ability to de-convolve neural responses to different experimental conditions. The materials for all experiments are available from OSF (https://osf.io/abjy9/).

**Figure 2:**
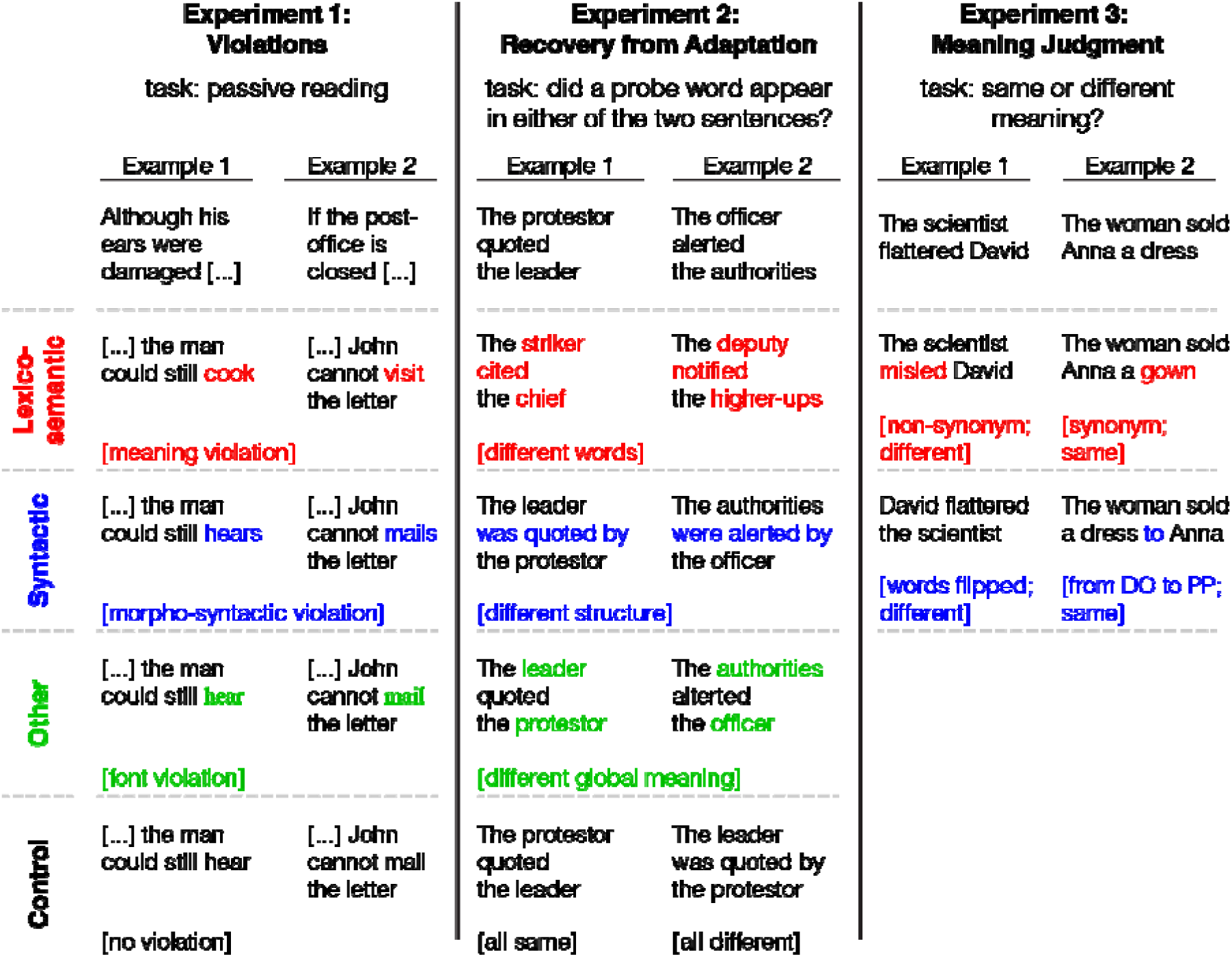
Sample stimuli for each condition in Experiments 1-3. Two examples are provided for each condition in each experiment. For Experiment 1, the top row shows the beginning of a sentence, and the next rows show different possible continuations. For Experiments 2-3, the top row shows one sentence from a pair, and the next rows show different possibilities for the other sentence in that pair. Red: Lexico-semantic condition; Blue: Syntactic condition; Green: other experimental conditions; Black: control condition. [NB1: For Experiment 1, the task was passive reading for the critical materials, but, as described in the text, a small number of (filler) trials contained a memory probe task. NB2: For Experiment 2, three versions of the same base item (corresponding to the Lexico-semantic, Syntactic, and Global meaning conditions) are presented for illustrative purposes. As detailed in the text, in the actual materials, distinct sets of base items were used for the three critical conditions in order to match the number of trials across conditions while avoiding sentence repetition.]

**Figure 3:**
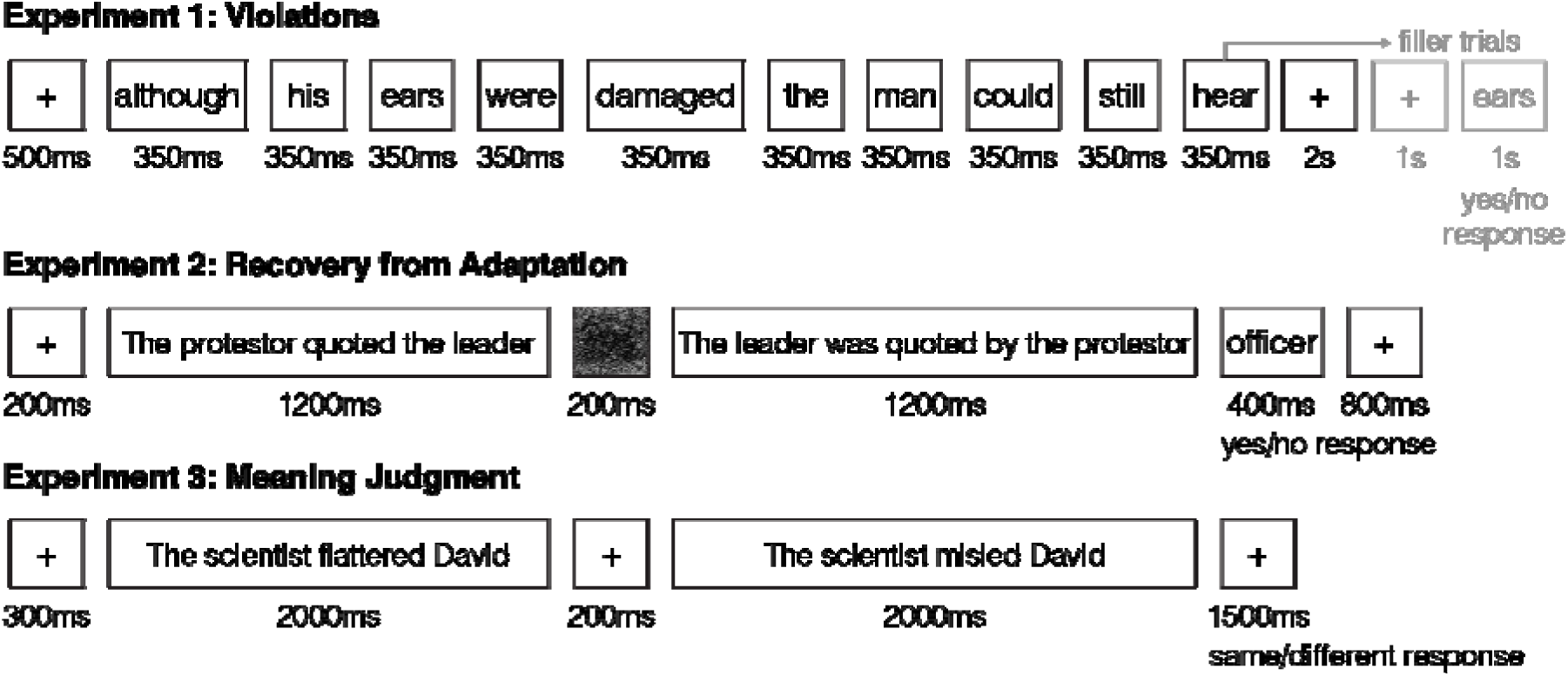
Trial structure for Experiments 1-3. One sample trial is shown for each experiment.

**Table 2.** Stimulus and Procedure details for each experiment.

|  | **Experiment 1:**<br>**Violations** | **Experiment 2:**<br>**Recovery from Adaptation** | **Experiment 3:**<br>**Meaning Judgment<sup>e</sup>** |
| --- | --- | --- | --- |
| Item description | 10-word sentence | Two transitive sentences A & B | Two sentences A & B |
| Base items | 240<sup>c</sup> | 144 × 3 sets | 80 |
| Versions per<br>base item<sup>a</sup> | 4<br>Lexico-semantic<br>Syntactic<br>Font<br>Control | 6<br>Critical<sup>d</sup>: A then B / B then A<br>Same: A then A / B then B<br>Different: X then A / Y then B<br>(X, Y: another item) | 4<br>Lexico-semantic, same<br>Lexico-semantic, different<br>Syntactic, same<br>Syntactic, different |
| Unique trials | 240 × 4 = 960 | (144 × 3) × 6 = 2592 | 80 × 4 = 320 |
| Experimental lists | 4<br>(240 trials each, 60<br>per version) | 6<br>(432 trials each, 144 per subset,<br>24 per version per subset) | 4<br>(80 trials each,<br>20 per version) |
| Trial breakdown<br>in a single list | 60 Lexico-semantic<br>60 Syntactic<br>60 Font<br>60 Control<br>(+10 fillers each) | 48 Critical Lexico-semantic<br>48 Critical Syntactic<br>48 Critical Global meaning<br>48 × 3 = 144 Same<br>48 × 3 = 144 Different | 40 Lexico-semantic<br>40 Syntactic |
| Runs per list | 5 (sometimes 4) | 6 (sometimes 5) | 2 |
| Trials per version<br>per run | 12<br>(+2 fillers) | 4<br>(24 per subset) | 10 |
| Trial length (s)<sup>b</sup> | 6 | 4 | 6 |
| Fixation per run (s) | 72 | 32 | 120 |
| Run duration (s) | 408 (6min 48s) | 320 (5min 20s) | 360 (6min) |
| No. condition orders | 10 | 6 | 8 |
| Subjects per list | 4-6 | 2-4 | 3-4 |

<sup>a</sup> See Figure 2.
<sup>b</sup> See Figure 3.
<sup>c</sup> Verbs did not repeat across items, except for “*practice*” and “*read*”, which were each used twice. 179 base items were adapted from Kuperberg et al. (2003).
<sup>d</sup> Each of the three sets of base items had a different critical condition: Lexico-semantic, Syntactic, or Global meaning (see Figure 2).
<sup>e</sup> Stimuli and procedure followed the design of Dapretto & Bookheimer (1999).

*Experiment 1: Lexico-semantic vs. (morpho-)syntactic violations.* Participants passively read stimuli, and their expectations were violated in several ways. The items were 10-word sentences, with four versions each (**Figure 2**): the critical verb (i) resulted in a lexico-semantic violation (stimuli that typically elicit an N400 component in ERP studies; see Kutas & Federmeier, 2011, for a review); (ii) resulted in a morpho-syntactic violation (stimuli that typically elicit a P600 component in ERP studies; e.g., Osterhout & Holcomb, 1992; Hagoort et al., 1993); (iii) contained no violations (control condition); or (iv) was presented in a different font (a low-level oddball violation, included as an additional, stricter, control condition). Lexico-semantic violations were created by shuffling the critical verbs across the base items. Syntactic violations were created by either omitting a required morpheme (30%) or adding an unnecessary morpheme (70%). (The reason we included a higher proportion of added compared to missing morphemes is because form-based error correction mechanisms are so robust during language comprehension that grammatical errors are often missed during proofreading (e.g., Schotter et al., 2014), and noticing missing elements seems harder than the extra ones.)

Overall, there were 240 items. They included 139 items with a sentence-final critical verb, taken (and sometimes slightly edited) from Kuperberg et al. (2003), as well as 61 additional items (to increase power) constructed in a similar manner. Further, to render the timing of violations less predictable, we adapted another 40 items from Kuperberg et al. such that the critical verb appeared before the final (10^th^) word: 6 items had the verb in each of the 3^rd^ through 8^th^ positions, and 4 items had it in the 9^th^ position. Critical verbs were not repeated across the 240 items, with two exceptions (“practice” and “read” were used twice each). For each participant, 10 additional sentences were included in each of the four conditions to serve as fillers. These fillers were followed by a memory-probe task (deciding whether the probe word appeared in the preceding sentence; **Figure 3**) to ensure that participants paid attention to the task; they were excluded from data analysis.

*Experiment 2: Recovery from adaptation to word meanings vs. syntactic structure.* Participants were asked to attentively read pairs of sequentially presented sentences and perform a memory probe task at the end of each pair (i.e., decide whether a probe word appeared in either of the two sentences). The sentences were simple transitive sentences consisting of an agent, a verb, and a patient. Because of constraints on these materials (as elaborated below), we constructed three sets of items (**Figure 2**): (i) sentence pairs that differed only in lexical items (but had the same syntactic structure and global meaning), created by replacing the verb and the agent and patient noun phrases with synonyms or words closely related in meaning; (ii) pairs that differed only in their syntactic structure (but had the same lexical items and global meaning), created by using the Active/Passive alternation; and (iii) pairs that differed only in the global meaning (but had the same lexical items and syntactic structure), created by switching the two noun phrases, leading to opposite thematic role assignment. The third set was included in order to examine sensitivity to overall propositional meaning and to probe combinatorial semantic processing. Overall, there were 432 items (144 per set).

In each set, each sentence pair {A,B} had six versions (**Table 2**): sentence A followed by sentence B, and sentence B followed by sentence A (“Critical” condition); sentence A followed by sentence A, and sentence B followed by sentence B (“Same” condition); and, finally, sentence A followed by a completely different sentence X (lexical items, syntactic structure, and global meaning were all different), and sentence B followed by a completely different sentence Y, where the pair {X,Y} was taken from another item (“Different” condition). Every sentence was used once in the Different condition of some other item. Therefore, within each of the three sets of items, every sentence appeared twice in each condition (Critical, Same, Different). Across the three sets, there were overall 5 experimental conditions: Critical Lexico-semantic, Critical Syntactic, Critical Global meaning, Same, and Different. In order to clearly mark the distinctness of the two identical sentences in the Same condition, trials across all conditions included a brief visual mask between the two sentences.

To keep the materials semantically diverse, items in the first two sets were constructed to be evenly distributed among three types of agent-patient relationships: (1) animate agent + inanimate patient; (2) animate agent + animate patient, where the relationship is biased so that one of the noun phrases is much more likely to be the agent (e.g., *The hit man killed the politician*); and (3) animate agent + animate patient, where the two nouns are similarly likely to be the agent (e.g., *The protestor quoted the leader*). By virtue of the manipulation of global meaning in the third set, all items had to be semantically reversible (i.e., of the third type).

*Experiment 3: Same-different meaning judgment on sentences that differ in word meanings vs. syntactic structure*

This design was adapted from Dapretto & Bookheimer (1999). Participants were asked to decide whether or not a pair of sequentially presented sentences had roughly the same meaning. The items were 80 sentence pairs, and each pair had four versions (**Figure 2**; **Table 2**): two versions in which the sentences differed in a single word (Lexico-semantic condition), replaced by either a synonym (Same meaning version) or a non-synonym (Different meaning version); and two versions (Syntactic condition) in which the sentences were either syntactic alternations differing in both structure and word order (Same meaning version), or in only structure / only word order (Different meaning version). Half of the items used the Active/Passive constructions (as in Dapretto & Bookheimer), and half – the Double Object (DO) / Prepositional Phrase Object (PP) constructions.

A number of features varied and were balanced across items (**Figure 2**). First, the construction was always the *same* across the two sentences in the Lexico-semantic condition (balanced between active and passive for the Active/Passive items, and between DO and PP for the DO/PP items). However, in the Syntactic condition, the construction was always *different* in the Same-meaning version because this is how the propositional meaning was preserved (again, balanced between active and passive for the Active/Passive items, and between DO and PP for the DO/PP items). For the Different-meaning version, the construction could either be the same (in which case the order of the two relevant nouns was switched) or different (in which case the order of the two relevant nouns was preserved). In cases where the construction differed across the two sentences, we balanced whether the first sentence was active *vs.* passive (for the Active/Passive items), or whether it was DO *vs.* PP (for the DO/PP items). The second feature that varied across the materials was whether the first-mentioned noun was a name or an occupation noun. All items contained one instance of each, with order of presentation balanced across stimuli. And third, for the Lexico-semantic condition, we varied how exactly the words in the second sentence in a pair differed from the words in the first. (This does not apply to the Syntactic condition because the content words were identical across the two sentences within each pair.) In particular, for the Active/Passive items, either the occupation noun or the verb could be replaced (by a synonym or a word with a different meaning); and for the DO/PP items, either the occupation noun or the direct object (inanimate) noun could be replaced.

### Data acquisition, preprocessing, and first-level modeling

#### Data acquisition

Whole-brain structural and functional data were collected on a whole-body 3 Tesla Siemens Trio scanner with a 32-channel head coil at the Athinoula A. Martinos Imaging Center at the McGovern Institute for Brain Research at MIT. T1-weighted structural images were collected in 176 axial slices with 1mm isotropic voxels (repetition time (TR) = 2,530ms; echo time (TE) = 3.48ms). Functional, blood oxygenation level-dependent (BOLD) data were acquired using an EPI sequence with a 90° flip angle and using GRAPPA with an acceleration factor of 2; the following parameters were used: thirty-one 4.4mm thick near-axial slices acquired in an interleaved order (with 10% distance factor), with an in-plane resolution of 2.1mm × 2.1mm, FoV in the phase encoding (A >> P) direction 200mm and matrix size 96 × 96 voxels, TR = 2000ms and TE = 30ms. The first 10s of each run were excluded to allow for steady state magnetization.

#### Preprocessing

Data preprocessing was carried out with SPM5 (using default parameters, unless specified otherwise) and supporting, custom MATLAB scripts. (Note that SPM was only used for preprocessing and basic modeling—aspects that have not changed much between versions; we used an older version of the SPM software because we have projects that use data collected over the last 15 years, and we want to keep preprocessing and first-level analysis the same across the 800+ subjects in our database, which are pooled in the analyses for many projects. For several datasets, we have directly compared the outputs of data preprocessed and modeled in SPM5 vs. SPM12, and the outputs are nearly identical.) Preprocessing of anatomical data included normalization into a common space (Montreal Neurological Institute (MNI) template) and resampling into 2mm isotropic voxels. Preprocessing of functional data included motion correction (realignment to the mean image of the first functional run using 2^nd^-degree b-spline interpolation), normalization (estimated for the mean image using trilinear interpolation), resampling into 2mm isotropic voxels, smoothing with a 4mm FWHM Gaussian filter and high-pass filtering at 200s.

#### Data modeling

For both the language localizer task and the critical tasks, a standard mass univariate analysis was performed in SPM5 whereby a general linear model (GLM) estimated the effect size of each condition in each experimental run. These effects were each modeled with a boxcar function (representing entire blocks/events) convolved with the canonical Hemodynamic Response Function (HRF). The model also included first-order temporal derivatives of these effects, as well as nuisance regressors representing entire experimental runs and offline-estimated motion parameters.

### Definition and validation of language-responsive functional regions of interest (fROIs)

For each participant (in each experiment), we defined a set of language-responsive functional ROIs using group-constrained, participant-specific localization (Fedorenko et al., 2010). In particular, each individual participant’s map for the sentences > nonwords contrast from the language localizer task was intersected with a set of six binary masks. These masks were derived from a probabilistic activation overlap map for the language localizer contrast in a large set of participants (n=220) using the watershed parcellation, as described in Fedorenko et al. (2010), and corresponded to relatively large areas within which most participants showed activity for the target contrast. These masks covered the fronto-temporal language network: three in the left frontal lobe falling within the IFG, its orbital portion, and the MFG, and three in the temporal and parietal cortex (**Figure 5**). Within each mask, a participant-specific language fROI was defined as the top 10% of voxels with the highest *t*-values for the localizer contrast. This top *n*% approach ensures that fROIs can be defined in every participant and that their sizes are the same across participants, allowing for generalizable results (e.g., Nieto-Castañón and Fedorenko, 2012).

Before examining the data from the critical experiments, we ensured that the language fROIs show the expected signature response (i.e., that the response is reliably greater to sentences than nonwords). To do so, we used an across-runs cross-validation procedure (e.g., Nieto-Castañón & Fedorenko, 2012), where one run of the localizer is used to define the fROIs, and the other run to estimate the responses, insuring independence (e.g., Kriegeskorte et al., 2009). As expected, and replicating prior work (e.g., Fedorenko et al., 2010; Fedorenko et al., 2011; Blank et al., 2016; Mahowald & Fedorenko, 2016), the language fROIs showed a robust sentences > nonwords effect (*t*s_(48)_>8.44; *p*s<0.0001), correcting for the number of regions (six) using the False Discovery Rate (FDR) correction (Benjamini & Yekutieli, 2001).

### Estimating the responses of the language fROIs to the conditions of the critical experiments

We estimated the responses in the language fROIs to the conditions of each critical experiment: the Control condition, Lexico-semantic violations, Syntactic violations, and Font violations in Experiment 1; the Same condition, Different condition, and three Critical conditions (differing in only lexical items, only syntactic structure, or only global meaning) in Experiment 2; and the Lexico-semantic and Syntactic conditions (each collapsed across same and different pairs) in Experiment 3. Statistical comparisons were performed on the estimated percent BOLD signal change (PSC) values in each region in each experiment.

We analyzed each experiment separately to allow for the possibility that syntax selectivity would be observed in just one of the experiments. Such a pattern could still be potentially informative and would be missed in an analysis that pools data from the three experiments. Furthermore, we examined each region separately, in line with our research question: whether *any* region within the language network is selective for syntactic over lexico-semantic processing.

#### Reality-check analyses: Testing for sensitivity to lexico-semantic and syntactic processing

First, we tested for basic sensitivity to lexico-semantic and syntactic manipulations. In each region, we used two-tailed paired-samples *t*-tests to compare the response to each critical (lexico-semantic and syntactic) condition to one or more control conditions. In Experiment 1, we compared the response to each critical violation condition (Lexico-semantic or Syntactic) against a) the Control condition with no violations, and, as an additional, stricter, baseline, b) the Font violation condition. In Experiment 2, we compared the Same and Different conditions to each other (a reality check to test for recovery from adaptation in the language regions when all the features of the sentence change), and then we compared each of the Critical conditions to the Same condition (where the same sentence is repeated exactly) to test for recovery from adaptation when the lexical items or the syntactic structure changes. The predictions are similar for Experiments 1 and 2: if a brain region is sensitive to lexical processing, the Lexico-semantic condition should elicit a stronger response than the control condition(s); similarly, if a brain region is sensitive to syntactic processing, the Syntactic condition should elicit a stronger response than the control condition(s). Finally, in Experiment 3, we compared the response to each condition (Lexico-semantic and Syntactic) against the low-level fixation baseline to ensure robust responses in the language regions to sentence comprehension. (Note that fixation was used here because, unlike in the other two experiments, there was no other baseline condition following Dapretto & Bookheimer’s (1999) design.) In each of these reality-check analyses, the results were FDR-corrected (Benjamini & Yekutieli, 2001) for the six regions.

#### Critical analyses (a): Directly comparing the Lexico-semantic and Syntactic conditions in the language fROIs

Next, we directly compared the Lexico-semantic and Syntactic conditions in each region in each experiment. If a brain region is selectively or preferentially engaged in syntactic processing, then we would expect to observe a reliably stronger response to the Syntactic condition than the Lexico-semantic condition. And if a brain region is selectively/preferentially engaged in lexico-semantic processing, we would expect to observe the opposite pattern. In these analyses, we report the results without a correction for the number of regions because we wanted to give syntactic selectivity—which we are arguing *against*—the best chance to reveal itself. (Of course, if an uncorrected *p*-value fails to reach significance, then the corrected one does, too.)

#### Critical analyses (b): Searching for voxels selective for syntactic (or lexico-semantic, for completeness) processing

One potential concern with the use of language fROIs is that each fROI is relatively large and the responses are averaged across voxels (e.g., Friston et al., 2006). Thus, fROI-based analyses may obscure underlying functional heterogeneity and potential selectivity for syntactic processing. For example, if a fROI contains a subset of voxels that show a stronger response to lexico-semantic than syntactic processing, and another subset of voxels that show a stronger response to syntactic than lexico-semantic processing, we may not detect a difference at the level of the fROI as a whole. To circumvent this concern, we supplemented the analyses of language fROIs, with analyses that i) use some of the data from each critical experiment to directly search for voxels— within the same broad masks encompassing the language network—that respond more strongly to syntactic than lexico-semantic processing (i.e., top 10% of voxels based on the Syntactic>Lexico-Semantic contrast), or vice versa (for completeness), and then ii) examine the replicability of this pattern of response in a left-out portion of the data. We performed this analysis for each of Experiments 1-3. If any (even non-contiguous) voxels with reliably stronger responses to syntactic processing exist anywhere within the fronto-temporal language network, this analysis should be able to detect them. For these analyses, we used one-tailed paired-samples *t*-tests because these hypotheses are directional. For example, when examining voxels that show stronger responses to syntactic than lexico-semantic processing to test whether this preference is replicable in left-out data, the critical contrast is Syntactic>Lexico-Semantic. As in the last set of analysis, these results were not corrected for the number of regions because we wanted to give syntactic selectivity the best chance to reveal itself.

## Results

### Behavioral results

Error rates and reaction times (RTs), for trials with a recorded response, in each of the three experiments are summarized in **Figure 4**. Performance on the memory probe task in the filler trials in Experiment 1 was close to ceiling (between 95.4% and 96.6% across conditions), with no reliable difference between the two critical—Lexico-semantic and Syntactic—conditions, in accuracies or RTs (*ts*_(21)_<1, *n.s.*). In Experiment 2, performance on the memory probe task varied between 72.6% and 95.7% across conditions. As expected, participants were faster and more accurate in the Same condition, where the same sentence was repeated, than in the Different condition, where the two sentences in the pair differed in all respects (*t*s_(13)_>15.1, *p*s<0.05). Furthermore, participants were faster and more accurate in the Syntactic condition than in the Lexico-semantic condition (*t*s(_13)_>3.55, *p*s<0.05), plausibly because the words were repeated between the two sentences in the pair in the Syntactic, but not Lexico-semantic condition. However, the difference was small, with high performance (>89.7%) in both conditions. Finally, in line with the behavioral results in Dapretto & Bookheimer (1999), in Experiment 3, performance on the meaning judgment task did not differ between the Lexico-semantic and Syntactic conditions, in accuracies or RTs (*ts*_(14)_<1, *n.s.*). In summary, in each of the three experiments, performance was high across conditions, with no reliable differences between the Lexico-semantic and Syntactic conditions in Experiments 1 and 3, and only a small difference in Experiment 2. We can thus proceed to examine neural differences between lexico-semantic and syntactic processing without worrying about those differences being driven by systematic differences in processing difficulty.

**Figure 4:**
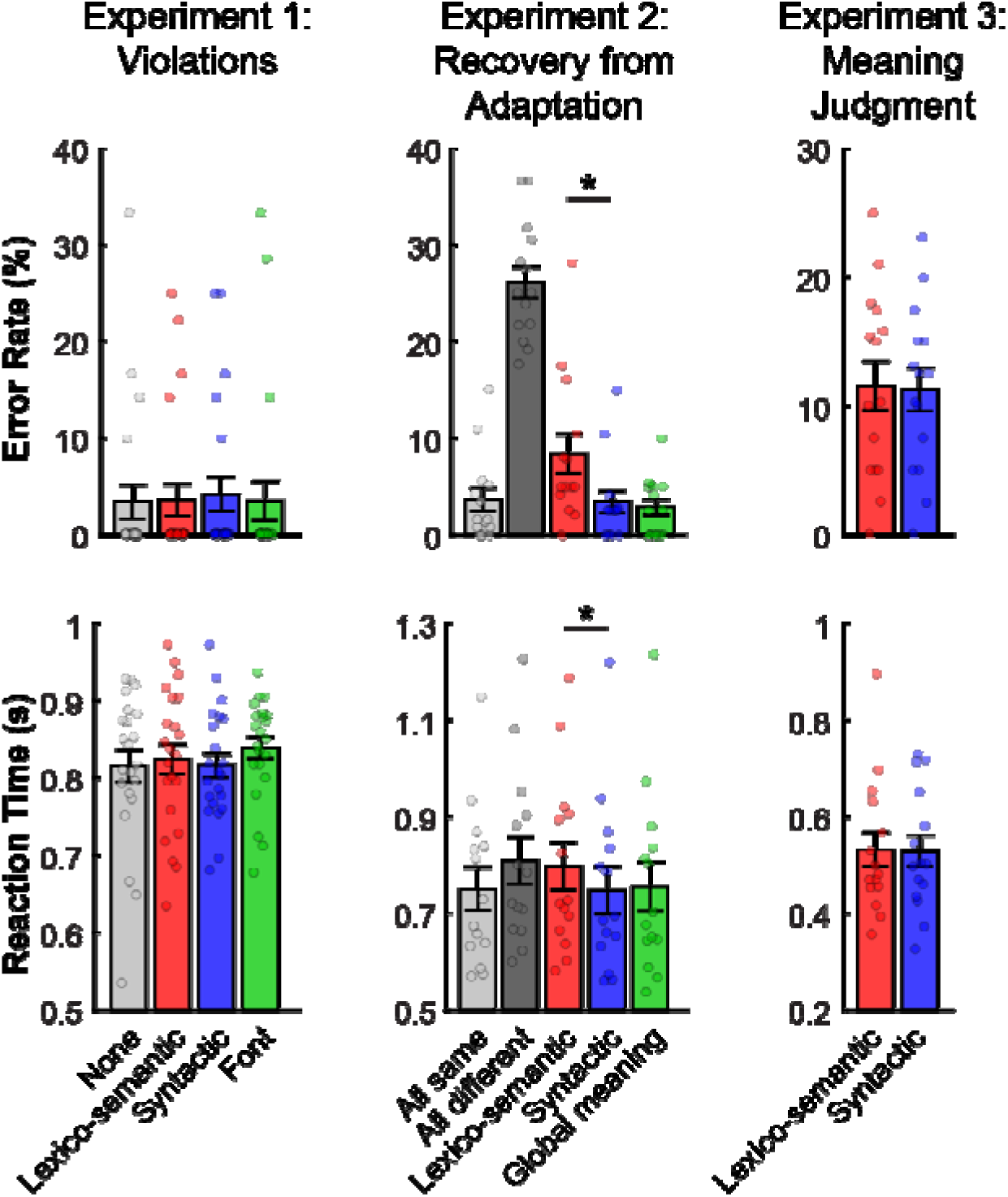
Summary of the behavioral results from Experiments 1-3. Each column shows the results from a different experiment: left – Experiment 1: Violations; middle – Experiment 2: Recovery from Adaptation; right – Experiment 3: Meaning Judgment. Top graphs: error rates; bottom graphs: reaction times. Here and in all subsequent figures, for each condition and each measure, dots correspond to individual participants; the bar shows the average across these participants, and the error bar shows standard error of the mean (across participants). Significant differences between the critical, Lexico-semantic (red) and Syntactic (blue), conditions are marked with *’s.

**Figure 5:**
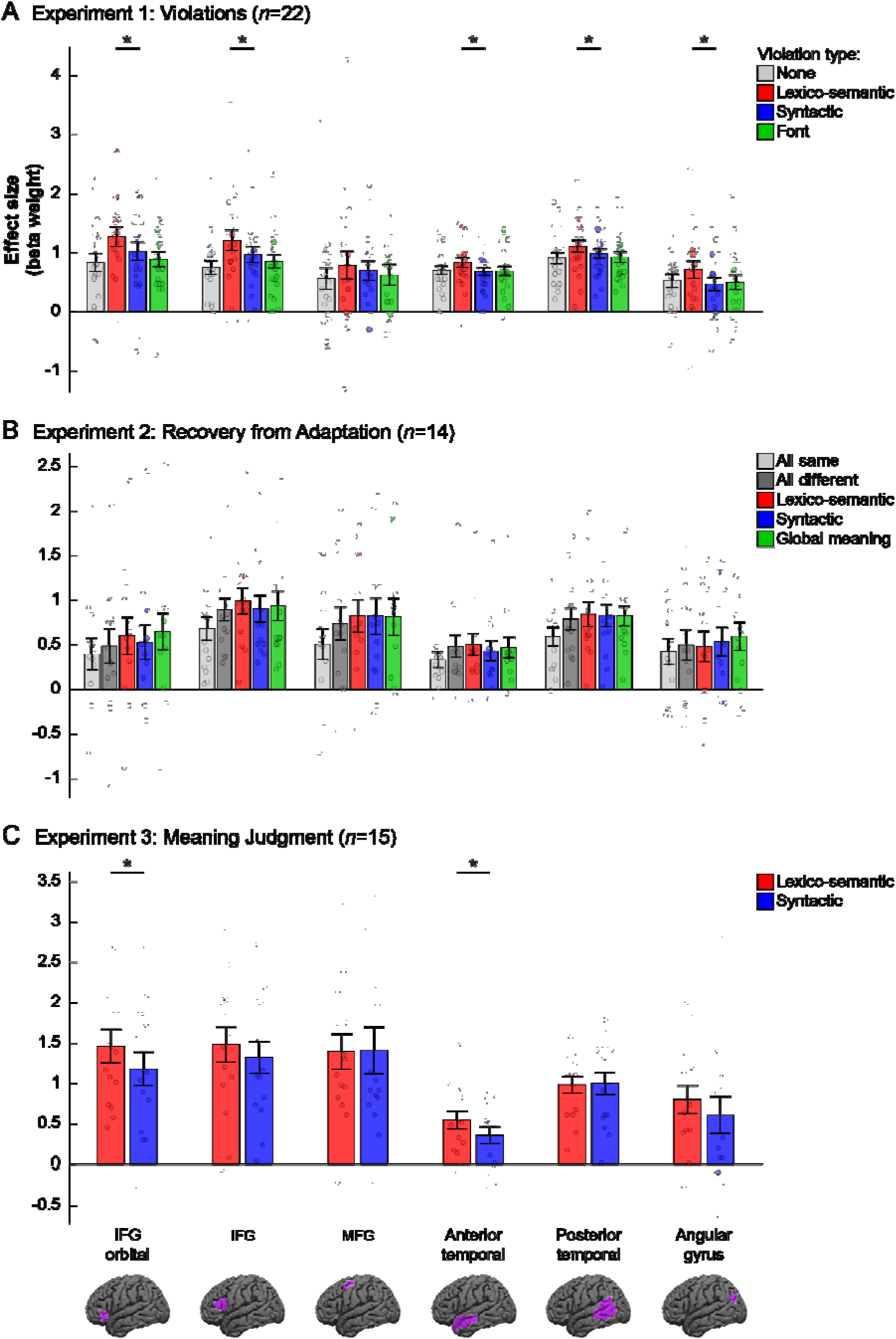
Responses in language fROIs to the conditions in Experiments 1-3. Responses (beta weights from a GLM) are measured as PSC relative to the fixation baseline. Each panel shows the results from a different experiment: (A) Experiment 1: Violations; (B) Experiment 2: Recovery from Adaptation; (C) Experiment 3: Meaning Judgment. Within each panel, each group of bars shows data from a different fROI (legend at the bottom). Significant differences between the critical, Lexico-semantic (red) and Syntactic (blue), conditions are marked with *’s. (The significance of the reality-check analyses that establish the sensitivity of the language fROIs to the lexico-semantic and syntactic manipulations relative to the control conditions is not marked; please see Results and Table 3.) Brain images at the bottom show the broad masks used to constrain the selection of the individual fROIs (these are not the fROIs themselves, which were defined using the sentences>nonwords contrast separately in each participant, as described in the text, and could thus vary within the borders of the masks depicted here).

### fMRI results

#### 1. Reality-check analyses

*Testing for sensitivity to lexico-semantic and syntactic processing.* The results for the three experiments are summarized in **Figure 5** and **Table 3**. In Experiment 1, the Lexico-semantic condition elicited a reliably stronger response than the Control (no violations) condition in each of six language fROIs (*t*s_(21)_>2.77, *p*s<0.05), and than the Font violations condition in five of the six fROIs (*t*s_(21)_>3.43, *p*s<0.05), with the MFG fROI not showing a significant effect. The Syntactic condition elicited a reliably stronger response than the Control condition in two language fROIs: IFGorb and IFG (*t*s_(21)_>2.84, *p*s<0.05). However, the Syntactic condition did not reliably differ from the Font violations condition in any of the fROIs (*t*s_(21)_<1.45, *n.s.*), suggesting that language regions are not recruited more strongly when people encounter syntactic violations than they are when people encounter low-level perceptually unexpected features in the linguistic input (see also Vissers et al., 2006; van de Meerendonk et al., 2011; see also Mollica et al., 2020, for extension to word-order violations). In Experiment 2, the Different condition—where the two sentences in a pair differed in lexical items, syntactic structure, and global meaning—elicited a reliably stronger response than the Same condition, where the two sentences in a pair were identical, in four language fROIs: IFG, MFG, AntTemp, and PostTemp (*t*s(13)>3.45, *p*s<0.05); the effect was not reliable in the IFGorb and AngG fROIs. Further, the Lexico-semantic condition elicited a response that was reliably stronger than the Same condition in all language fROIs, except for the AngG fROI (*t*s_(13)_>2.71, *p*s<0.05), and similarly, the Syntactic condition elicited a response that was reliably stronger than the Same condition in all language fROIs, except for the AngG fROI (*t*s_(13)_>2.33, *p*s<0.05)2 . Finally, in Experiment 3, both experimental conditions elicited responses that were reliably above the fixation baseline in all six fROIs (Lexico-semantic: *t*s_(14)_>4.58, *p*s<0.05; Syntactic: *t*s_(14)_>2.66, *p*s<0.05).

**Table 3.** Responses of language fROIs in Experiments 1-3: mean PSC with standard error (by participants), effect size (Cohen’s *d*), *t*-value, and *p*-value.^a^

| fROI<br>(size) | Exp. 1: violations |  |  |  |  | Exp. 2: adaptation recovery |  |  |  | Exp. 3: meaning judgments |  |  |
| --- | --- | --- | --- | --- | --- | --- | --- | --- | --- | --- | --- | --- |
|  | LexSem | Synt | LexSem | Synt | LexSem | Diff | LexSem | Synt | LexSem | LexSem | Synt | LexSem |
|  | vs.<br>Control | vs.<br>Control | vs.<br>Font | vs.<br>Font | vs.<br>Synt | vs.<br>Same | vs.<br>Same | vs.<br>Same | vs.<br>Synt | vs.<br>Baseline | vs.<br>Baseline | vs.<br>Synt |
| IFGorb<br>(37 voxels) | 0.44<br>($\pm 0.07$)<br>$d=1.37$<br>$t=6.40$<br>$p<10^{-5}$ | 0.19<br>($\pm 0.07$)<br>$d=0.61$<br>$t=2.85$<br>$p=.0096$ | 0.38<br>($\pm 0.08$)<br>$d=0.97$<br>$t=4.54$<br>$p=.0002$ | 0.13<br>($\pm 0.09$)<br>$d=0.31$<br>$t=1.44$<br>$p=.16$ | 0.25<br>($\pm 0.08$)<br>$d=0.64$<br>$t=3.01$<br>$p=.0067$ | 0.09<br>($\pm 0.05$)<br>$d=0.52$<br>$t=1.93$<br>$p=.075$ | 0.21<br>($\pm 0.08$)<br>$d=0.73$<br>$t=2.72$<br>$p=.18$ | 0.13<br>($\pm 0.04$)<br>$d=0.83$<br>$t=3.10$<br>$p=.0084$ | 0.07<br>($\pm 0.07$)<br>$d=0.28$<br>$t=1.05$<br>$p=0.31$ | 1.46<br>($\pm 0.21$)<br>$d=1.80$<br>$t=6.96$<br>$p<10^{-5}$ | 1.18<br>($\pm 0.21$)<br>$d=1.46$<br>$t=5.67$<br>$p<10^{-4}$ | 0.28<br>($\pm 0.12$)<br>$d=0.59$<br>$t=2.28$<br>$p=.039$ |
| IFG<br>(74 voxels) | 0.46<br>($\pm 0.08$)<br>$d=1.17$<br>$t=5.51$<br>$p<10^{-4}$ | 0.22<br>($\pm 0.06$)<br>$d=0.72$<br>$t=3.37$<br>$p=.0029$ | 0.35<br>($\pm 0.09$)<br>$d=0.88$<br>$t=4.12$<br>$p=.0005$ | 0.12<br>($\pm 0.09$)<br>$d=0.28$<br>$t=1.30$<br>$p=.21$ | 0.24<br>($\pm 0.08$)<br>$d=0.64$<br>$t=3.02$<br>$p=.0065$ | 0.21<br>($\pm 0.05$)<br>$d=1.08$<br>$t=4.02$<br>$p=.0014$ | 0.31<br>($\pm 0.08$)<br>$d=1.02$<br>$t=3.80$<br>$p=.0022$ | 0.22<br>($\pm 0.08$)<br>$d=0.74$<br>$t=2.77$<br>$p=.016$ | 0.09<br>($\pm 0.08$)<br>$d=0.29$<br>$t=1.09$<br>$p=0.29$ | 1.48<br>($\pm 0.21$)<br>$d=1.78$<br>$t=6.89$<br>$p<10^{-5}$ | 1.32<br>($\pm 0.20$)<br>$d=1.72$<br>$t=6.68$<br>$p<10^{-4}$ | 0.16<br>($\pm 0.12$)<br>$d=0.34$<br>$t=1.30$<br>$p=.21$ |
| MFG<br>(46 voxels) | 0.22<br>($\pm 0.08$)<br>$d=0.59$<br>$t=2.78$<br>$p=.011$ | 0.14<br>($\pm 0.09$)<br>$d=0.33$<br>$t=1.56$<br>$p=.13$ | 0.16<br>($\pm 0.09$)<br>$d=0.39$<br>$t=1.81$<br>$p=.084$ | 0.08<br>($\pm 0.09$)<br>$d=0.17$<br>$t=0.82$<br>$p=.042$ | 0.09<br>($\pm 0.12$)<br>$d=0.16$<br>$t=0.74$<br>$p=0.47$ | 0.23<br>($\pm 0.07$)<br>$d=0.93$<br>$t=3.47$<br>$p=.0042$ | 0.32<br>($\pm 0.10$)<br>$d=0.90$<br>$t=3.35$<br>$p=.0052$ | 0.32<br>($\pm 0.09$)<br>$d=0.92$<br>$t=3.46$<br>$p=.0043$ | 0.00<br>($\pm 0.08$)<br>$d=0.01$<br>$t=0.04$<br>$p=0.97$ | 1.39<br>($\pm 0.22$)<br>$d=1.64$<br>$t=6.36$<br>$p<10^{-4}$ | 1.41<br>($\pm 0.29$)<br>$d=1.25$<br>$t=4.85$<br>$p=.0002$ | -0.01<br>($\pm 0.12$)<br>$d=-0.03$<br>$t=-0.12$<br>$p=.90$ |
| AntTemp<br>(162 voxels) | 0.13<br>($\pm 0.03$)<br>$d=0.83$<br>$t=3.90$<br>$p=.0008$ | -0.02<br>($\pm 0.02$)<br>$d=-0.15$<br>$t=-0.71$<br>$p=.49$ | 0.14<br>($\pm 0.04$)<br>$d=0.85$<br>$t=3.99$<br>$p=.0007$ | -0.01<br>($\pm 0.04$)<br>$d=-0.04$<br>$t=-0.18$<br>$p=0.86$ | 0.15<br>($\pm 0.04$)<br>$d=0.87$<br>$t=4.10$<br>$p=.0005$ | 0.15<br>($\pm 0.04$)<br>$d=0.94$<br>$t=3.51$<br>$p<.0038$ | 0.17<br>($\pm 0.05$)<br>$d=0.95$<br>$t=3.56$<br>$p=.0035$ | 0.10<br>($\pm 0.04$)<br>$d=0.62$<br>$t=2.34$<br>$p=.036$ | 0.07<br>($\pm 0.05$)<br>$d=0.38$<br>$t=1.43$<br>$p=0.17$ | 0.54<br>($\pm 0.11$)<br>$d=1.26$<br>$t=4.87$<br>$p=.0002$ | 0.36<br>($\pm 0.11$)<br>$d=0.87$<br>$t=3.36$<br>$p=.0047$ | 0.19<br>($\pm 0.06$)<br>$d=0.81$<br>$t=3.14$<br>$p=.0072$ |
| PostTemp<br>(294 voxels) | 0.20<br>($\pm 0.04$)<br>$d=1.16$<br>$t=5.42$<br>$p<10^{-4}$ | 0.07<br>($\pm 0.04$)<br>$d=0.38$<br>$t=1.80$<br>$p=.086$ | 0.18<br>($\pm 0.05$)<br>$d=0.80$<br>$t=3.76$<br>$p=.0011$ | 0.06<br>($\pm 0.07$)<br>$d=0.18$<br>$t=0.85$<br>$p=.40$ | 0.12<br>($\pm 0.06$)<br>$d=0.45$<br>$t=2.13$<br>$p=.045$ | 0.20<br>($\pm 0.04$)<br>$d=1.22$<br>$t=4.58$<br>$p=.0005$ | 0.25<br>($\pm 0.06$)<br>$d=1.05$<br>$t=3.91$<br>$p=.0018$ | 0.24<br>($\pm 0.06$)<br>$d=1.07$<br>$t=3.99$<br>$p=.0015$ | 0.01<br>($\pm 0.07$)<br>$d=0.05$<br>$t=0.20$<br>$p=0.84$ | 0.98<br>($\pm 0.10$)<br>$d=2.43$<br>$t=9.43$<br>$p<10^{-6}$ | 1.00<br>($\pm 0.14$)<br>$d=1.85$<br>$t=7.16$<br>$p<10^{-5}$ | -0.02<br>($\pm 0.08$)<br>$d=-0.06$<br>$t=-0.22$<br>$p=.83$ |
| AngG<br>(64 voxels) | 0.19<br>($\pm 0.06$)<br>$d=0.73$<br>$t=3.40$<br>$p=.0027$ | -0.06<br>($\pm 0.04$)<br>$d=-0.30$<br>$t=-1.42$<br>$p=.17$ | 0.22<br>($\pm 0.06$)<br>$d=0.73$<br>$t=3.44$<br>$p=.0024$ | -0.03<br>($\pm 0.06$)<br>$d=-0.12$<br>$t=-0.58$<br>$p=.57$ | 0.25<br>($\pm 0.06$)<br>$d=0.95$<br>$t=4.48$<br>$p=.0002$ | 0.07<br>($\pm 0.05$)<br>$d=0.43$<br>$t=1.59$<br>$p=.13$ | 0.06<br>($\pm 0.06$)<br>$d=0.28$<br>$t=1.04$<br>$p=.32$ | 0.11<br>($\pm 0.05$)<br>$d=0.58$<br>$t=2.16$<br>$p=.054$ | -0.05<br>($\pm 0.07$)<br>$d=-0.21$<br>$t=-0.79$<br>$p=0.44$ | 0.80<br>($\pm 0.17$)<br>$d=1.18$<br>$t=4.59$<br>$p<.0004$ | 0.61<br>($\pm 0.23$)<br>$d=0.69$<br>$t=2.67$<br>$p=.018$ | 0.19<br>($\pm 0.14$)<br>$d=0.34$<br>$t=1.33$<br>$p=.20$ |

<sup>a</sup> non-significant effects at $\alpha=0.05$ (FDR-corrected for the number of regions) are shaded in grey; marginal effects ( $0.05<p<0.1$ ) are shaded in light grey.

#### 2. Critical analyses (a): Directly comparing the Lexico-semantic and Syntactic conditions in the language fROIs

The results for the three experiments are summarized in **Figure 5** and **Table 3**. In Experiment 1, a direct comparison between the Lexico-semantic and Syntactic conditions revealed reliably stronger responses to the Lexico-semantic condition in all language fROIs except for the MFG fROI (*t*s_(21)_>2.12, *p*s<0.05). In Experiment 2, no language fROI showed reliably stronger recovery from adaptation in the Lexico-semantic than Syntactic condition or vice versa (*t*s_(13)_<1.44, *n.s.*). Finally, in Experiment 3, we observed a stronger response to the Lexico-semantic than Syntactic condition in two language fROIs—IFGorb and AntTemp (*ts*_(14)_>2.27, *ps*<0.05). No language fROI showed the opposite pattern.

#### 3. Critical analyses (b): Searching for voxels selective for syntactic (or lexico-semantic, for completeness) processing

The results for the three experiments are summarized in **Figures 6** and **7** and **Table 4**. In Experiment 1, when we defined the individual fROIs by the Syntactic>Lexico-semantic contrast, we did not find a replicable (across runs) Syntactic>Lexico-semantic effect within any of the masks. In contrast, when we defined the fROIs by the Lexico-semantic>Syntactic contrast, we found a replicable Lexico-semantic>Syntactic effect within all the language masks, except for the MFG mask (*t*s_(21)_>2.75, *p*s<0.05), consistent with our finding of stronger responses for the Lexico-semantic than Syntactic condition in the fROIs that were defined by the language localizer contrast.

**Figure 6:**
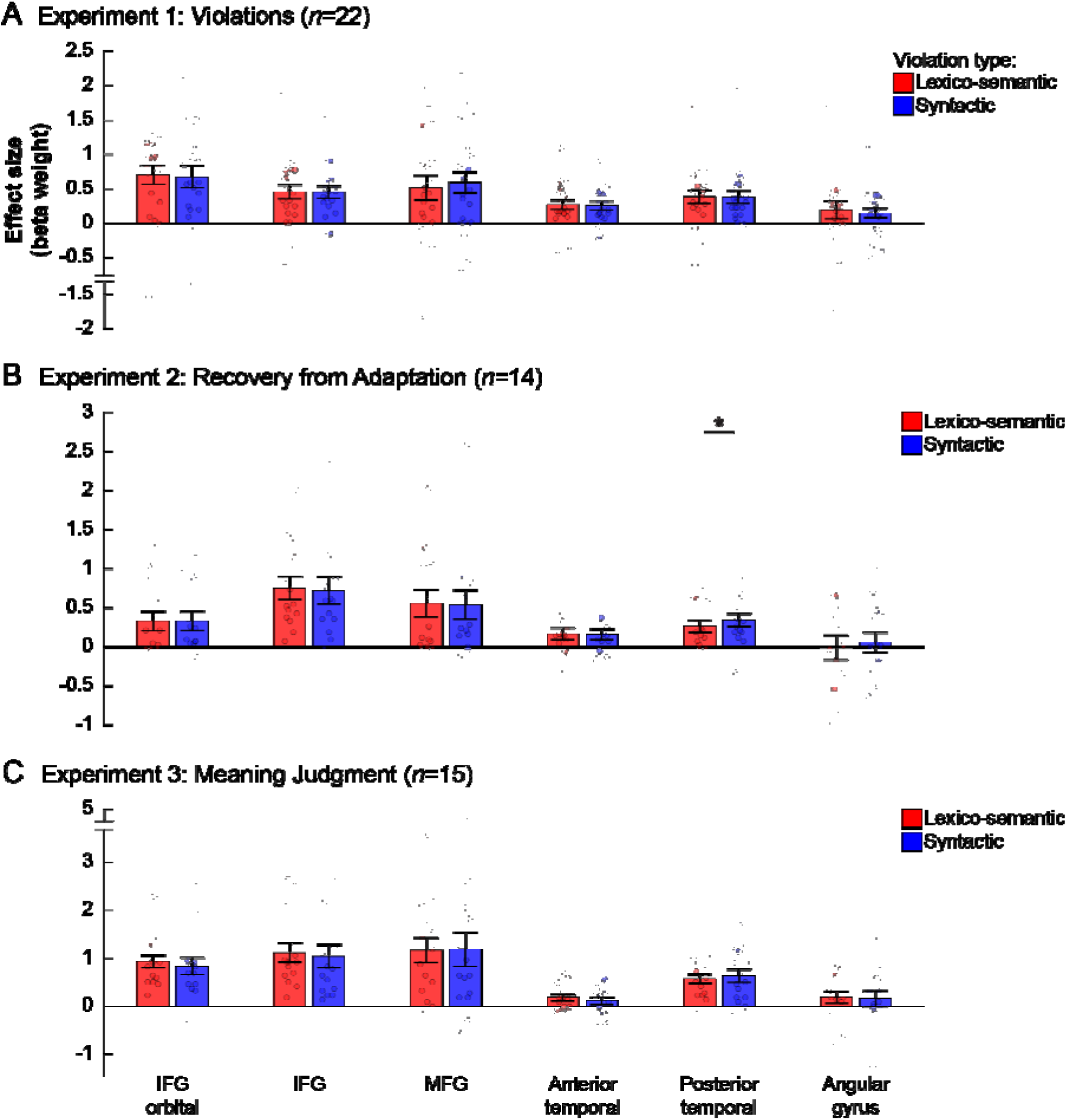
Responses in fROIs defined by the Syntactic>Lexico-semantic contrast to the critical conditions in Experiments 1-3. Participant-specific fROIs were defined, within the borders of each mask (Figure 5), as the top 10% of voxels showing the strongest Syntactic>Lexico-semantic contrast effect in the corresponding experiment. These fROIs were defined based on half the data from that experiment, and then the other (independent) half were used to estimate the effect size of this same contrast (i.e., estimate the replicability of the contrast effect). Conventions are the same as in Figure 5, with one exception: in panels A and C, parts of the *y*-axis at the top or bottom have been cut out (marked by two parallel horizontal tick marks) in order to stretch the bars more and accentuate differences across conditions when those appeared. In these visually edited cases, distance between the most extreme 1-2 data points and their corresponding bars are not at scale. Those data points are colored in gray. Differences between the Lexico-semantic (red) and Syntactic (blue) conditions are marked with *’s.

**Figure 7:**
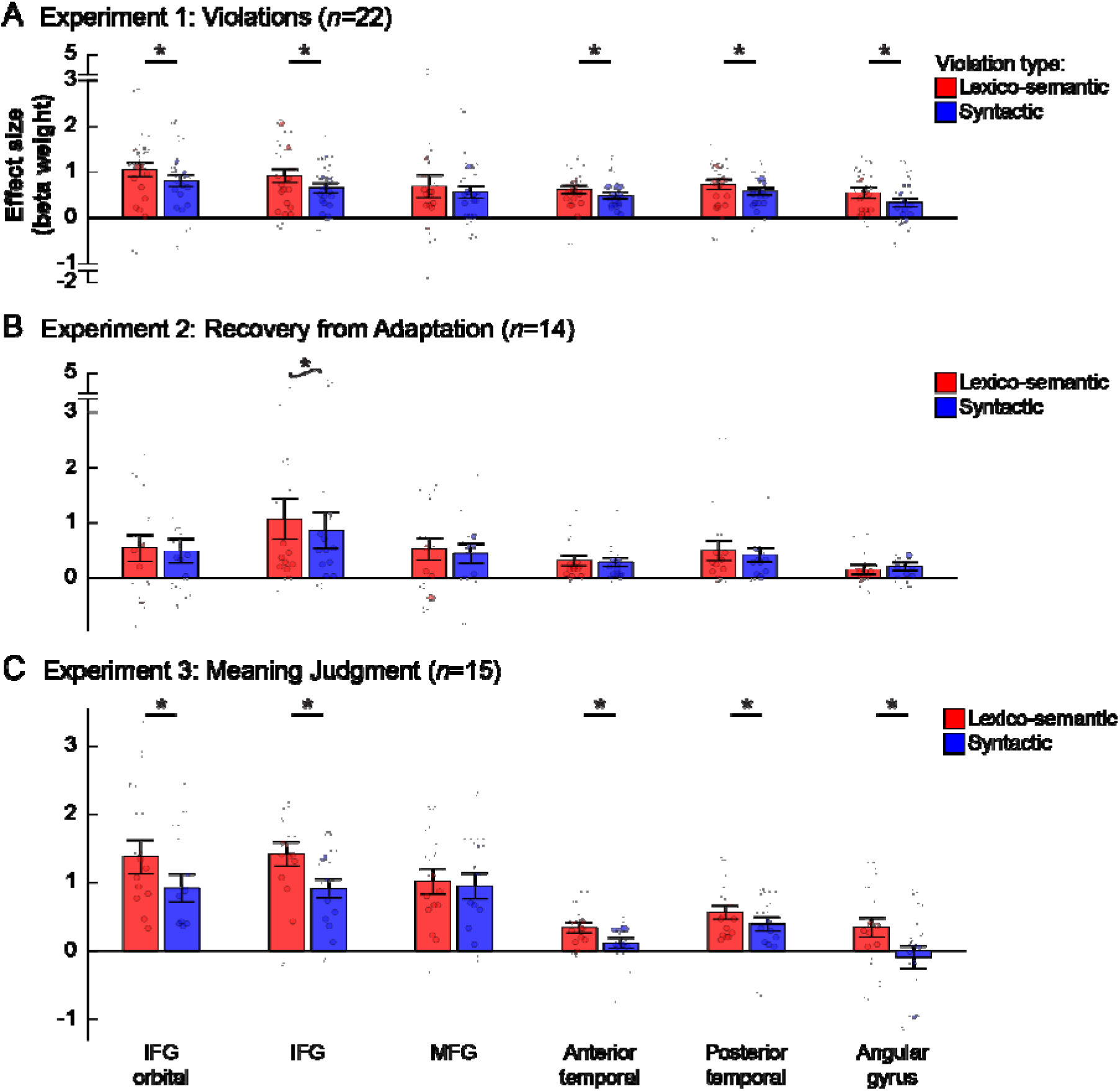
Responses in fROIs defined by the Lexico-semantic>Syntactic contrast to the critical conditions in Experiments 1-3. This figure depicts data from a parallel analysis to that depicted in Figure 6; here, participant-specific fROIs were defined as the top 10% of voxels showing the strongest Lexico-semantic>Syntactic contrast effect in the corresponding experiment, and the size of this contrast was then estimated in an independent part of the data (this is the opposite contrast to the one used in Figure 6). Conventions are the same as in Figures 5,6, with the addition of the following: non-significant effects with *p*<0.10 are marked with *s above tildes.

**Table 4.**
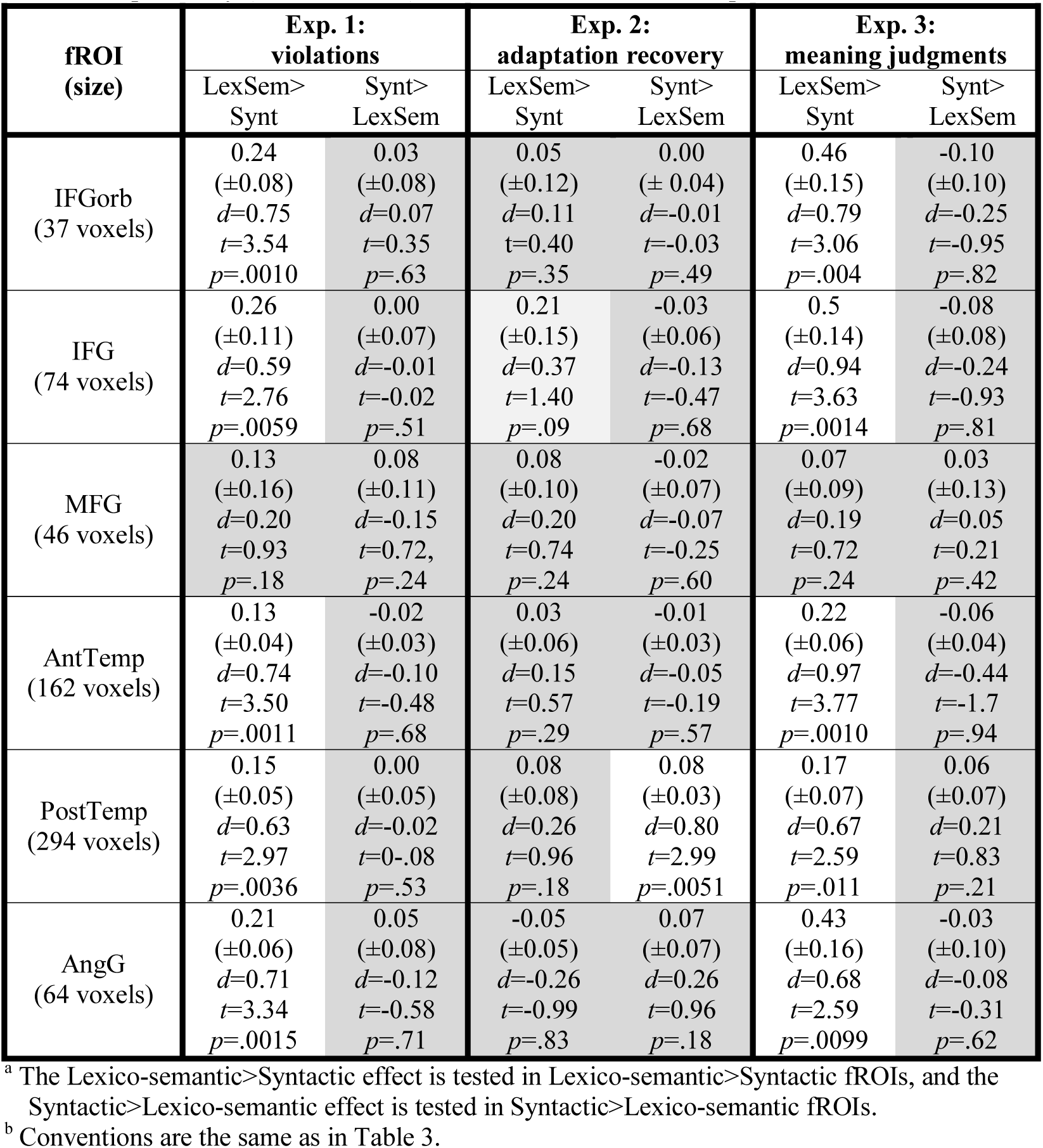
Replicability (in left-out data) of the critical contrasts in Experiments 1-3.^a,b^

In Experiment 2, when we defined the individual fROIs by the Syntactic>Lexico-semantic contrast, we found a slightly higher response to the Syntactic than Lexico-semantic condition within the PostTemp mask (*t*_(14)_=1.84, *p*<0.05). This effect would not survive correction for the number of regions. And even if, with a larger number of participants, the statistical robustness of the Syntactic>Lexico-semantic effect increased, the difference between the two conditions would remain small. When we defined the fROIs by the Lexico-semantic>Syntactic contrast, we did not find a replicable Lexico-semantic>Syntactic effect in any of the language masks, in line with the two critical conditions not eliciting differential responses in the fROIs defined by the language localizer contrast in this experiment.

Finally, in Experiment 3, as in Experiment 1, when we defined the individual fROIs by the Syntactic>Lexico-semantic contrast, we did not find a replicable Syntactic>Lexico-semantic effect within any of the masks; but when we defined the fROIs by the Lexico-semantic>Syntactic contrast, we found a replicable Lexico-semantic>Syntactic effect within all the language masks, except for the MFG mask (*t*s_(14)_>2.14, *p*s<0.05).

## Discussion

### Summary of the findings and their relationship to earlier proposals of the neural architecture of language

Across three fMRI experiments that used paradigms from the prior language literature but relied on a more sensitive and rigorous analytic approach—functional localization at the individual-subject level (Saxe et al., 2006; Fedorenko et al., 2010; Nieto-Castañón & Fedorenko, 2012)—we searched for syntactic selectivity within the fronto-temporal language network, asking whether any part of this network responds reliably more strongly to syntactic than lexico-semantic processing. No such selectivity was found. In particular, no language fROI, defined with a robust language localizer contrast (Fedorenko et al., 2010), showed stronger responses to syntactic processing than lexico-semantic processing in any experiment (**Figure 5**). Even sets of voxels defined by their stronger responses to syntactic than lexico-semantic conditions using half of the data did not show a replicable Syntactic>Lexico-semantic effect in the other half of the data (**Figure 6**), with one exception of a small difference between the Syntactic and Lexico-semantic conditions in one experiment that did not survive correction for the number of regions.

Importantly, this failure to uncover any syntactically-selective regions/voxels within the language network is not due to lack of power. Specifically, the lack of syntactic selectivity stands in sharp contrast to a) sensitivity to both lexico-semantic and syntactic manipulations—in at least a subset of the language fROIs—relative to the control conditions, where present (in Experiments 1 and 2) (**Figure 5**, **Table 3**); and b) stronger responses to lexico-semantic than syntactic conditions, replicable across some experiments (Experiments 1 and 3) and analyses (**Figures 5** and **7**, **Tables 3-4**). And although the lack of syntactic selectivity in the current study appears to run contrary to earlier brain imaging reports of dissociable effects of syntactic and semantic processing (e.g., Dapretto & Bookheimer, 1999; Embick et al., 2000; Kuperberg et al., 2000; Ni et al., 2000; Newman et al., 2001; Kuperberg et al., 2003; Noppeney & Price, 2004; Cooke et al., 2006; Friederici et al., 2010; Glaser et al., 2013; Schell et al., 2017, among others), none of those earlier studies had compellingly established selectivity for syntactic processing in a robust (across multiple sets of participants/materials) and generalizable (across diverse manipulations that aim to isolate the same cognitive process, and relative to diverse control conditions) way, as discussed in the Introduction.

The current results—along with earlier results from paradigms that have varied the presence/absence of lexico-semantic vs. syntactic information in the linguistic signal (e.g., Fedorenko et al., 2010; 2012a; 2016; Bedny et al., 2011; Pallier et al., 2011; Bautista & Wilson, 2016)—pose a challenge for proposals of the neural architecture of language that postulate syntax- or combinatorics-selective brain regions (e.g., Grodzinsky & Santi, 2008; Baggio and Hagoort, 2011; Friederici, 2011, 2012; Tyler et al., 2011; Duffau et al., 2014; Ullman, 2016; Matchin & Hickok, 2019; Pylkkanen, 2019). Across such proposals (see next section), a syntax-/combinatorics-selective component is argued to not support the storage and processing of individual word meanings, as illustrated in the architectures in **Figure 1a-d**. In contrast to these proposals, it appears that any brain region / set of voxels within the language network that shows sensitivity to syntactic manipulations also shows sensitivity to manipulations targeting the processing of individual word meanings.

Not all cognitive neuroscience proposals of the language architecture have postulate a distinction between syntactic and semantic representations/processing, or between combinatorial processing and stored knowledge representations (e.g., Bornkessel-Schlesewsky & Schlesewsky, 2009; Bornkessel-Schlesewsky et al., 2015). For example, Bornkessel-Schlesewsky et al. (2015) have suggested that the language network, as a whole, supports composition—combining smaller linguistic units into larger ones—in the service of meaning extraction (see also Mollica et al., 2020, for further empirical support of this idea). The current results align well with these proposals.

### The bias of the language network toward lexico-semantic processing

In two of our experiments, lexico-semantic conditions elicited numerically, and sometimes reliably, stronger responses than syntactic conditions in many of the language fROIs. Furthermore, in Experiment 1, unlike the lexico-semantic violation condition, the syntactic violation condition did not elicit a response that was higher than a low-level (font violation) condition in the language regions. These overall stronger responses to semantic than syntactic conditions are in line with two prior findings. First, using multivariate analyses, we have previously found that lexico-semantic information is represented more robustly than syntactic information in the language system (Fedorenko et al., 2012a). In particular, pairs of conditions that differ in whether or not they contain lexico-semantic information (e.g., sentences vs. Jabberwocky sentences, or lists of words vs. lists of nonwords) are more robustly dissociable in the fine-grained patterns of activity than pairs of conditions that differ in whether or not they are structured (e.g., sentences vs. lists of words, or Jabberwocky sentences vs. lists of nonwords). And second, in ECoG, we observed reliably stronger responses to conditions that only contain lexico-semantic information (word lists) than conditions that only contain syntactic information (Jabberwocky) in many language-responsive electrodes (Fedorenko et al., 2016), but no electrodes showed the opposite pattern. Along with the current study, these results demonstrate that the magnitude and spatial organization of responses in the human language network are determined more by meaning than structure.

This bias toward lexico-semantic processing, fits with the view that the goal of language is communication, i.e., the transfer of *meanings* across minds (e.g., Hurford, 1998, 2007; Goldberg, 2006; Jackendoff, 2011; Kirby et al., 2015; Gibson et al., 2018; Hahn et al., 2020), and with the fact that most of our knowledge of language has to do with lexical semantics (word meanings), with only a small number of bits needed to store all of our syntactic knowledge (Mollica and Piantadosi, 2019). And it is not consistent with syntax-centric views of language (e.g., Chomsky and DiNozzi, 1972; Pinker, 1995; Hauser et al., 2002; Friederici et al., 2006; Berwick et al., 2013; Friederici, 2018).

One implication of these, and earlier behavioral, results discussed in the Introduction is that artificial grammar learning and processing paradigms (e.g., Reber, 1967)—where structured sequences of meaningless units (e.g., syllables) are used in an attempt to approximate human syntax (e.g., Friederici et al., 2006; Petersson et al., 2010; Wang et al., 2015)—have limited utility for understanding human language, given that syntactic representations and processing seem to be inextricably linked with representations of linguistic meaning (see also Fedor et al., 2012).

### Limitations of scope

It is worth acknowledging the limitations of the current study. ***First***, perhaps some *cells / circuits / cortical-layer-specific areas* are selective for (some aspect) of syntactic/combinatorial processing (e.g., the architecture in **Figure 1e**), but we are not able to detect this selectivity due to the relatively coarse spatial resolution of our method. This possibility is hard to rule out without resorting to approaches like single-cell recordings (e.g., Engel et al., 2005; Mukamel & Fried, 2012) or laminar imaging (e.g., Norris & Polimeni, 2019). However, it is worth noting that at least for the paradigm that varies the presence/absence of lexico-semantic vs. syntactic information in the linguistic signal, the results from an ECoG study (Fedorenko et al., 2016)—where the spatial resolution is substantially higher than in fMRI—closely mirrored the fMRI results (e.g., Fedorenko et al., 2010).

Another possibility is that no single brain region is selective for syntactic/combinatorial processing, but inter-region *interaction/synchronization*—perhaps restricted to particular frequency bands (e.g., Giraud & Poeppel, 2012; Meyer, 2017; Martin & Doumas, 2019)—is critical for syntactic structure building. Some have argued that the arcuate / superior longitudinal fasciculus—the dorsal tract that connects posterior temporal and inferior frontal language areas—is critical for syntactic processing (e.g., Friederici, 2009; Brauer et al., 2011; Papoutsi et al., 2011; Wilson et al., 2011). However, the *selectivity* of this tract for syntactic/combinatorial processing is unclear, as it has also been implicated in non-syntactic computations, including, most commonly, articulation (e.g., Duffau et al., 2003; Hickok & Poeppel, 2007; Rauscheker & Scott, 2009), but also aspects of semantic processing (e.g., Glasser & Rillings, 2008). Thus, we would argue that, at present, no unequivocal evidence of syntax selectivity exists for inter-regional connections either.

***Second***, our research question focused on the “core” fronto-temporal language network, consisting of regions on the lateral surfaces of left frontal and left temporal cortex (e.g., Fedorenko & Thompson-Schill, 2014). Some areas *outside of this network’s boundaries* have been implicated in syntactic processing, including, for example, parts of the basal ganglia (e.g., Ullman, 2001, 2004; cf. Grossman et al., 2002; Longworth et al., 2005) or the cerebellum (see Marien et al., 2014 for a review). However, we would argue that, similar to the cortical language regions, noone has compellingly established selectivity for syntactic processing for any brain region outside of the core language network. Of relevance to this question is a claim that a region residing in or around Broca’s area supports abstract hierarchical structure processing across domains, including language, arithmetic, music, and action observation/planning (e.g., Koechlin & Jubault, 2006; Tettamanti & Weniger, 2006; Fadiga et al., 2009; Fitch & Martins, 2014). This claim does not find empirical support. In particular, the part of Broca’s area that responds to the presence of structure in language (e.g., showing stronger responses to structured linguistic stimuli, like sentences, than to lists of unconnected words) is highly selective for language relative to non-linguistic tasks, including ones that involve hierarchical structure and/or recursion, like arithmetic and music (e.g., Fedorenko et al., 2011; Monti et al., 2012; see Fedorenko & Varley, 2016, for a review). These results make sense given the strong links between linguistic structure and meaning discussed in this manuscript: in other words, given what we now know about linguistic syntax, the idea that it would be supported by a mechanism that is not sensitive to the nature of the representation does not seem tenable.

One possible explanation for some of the findings that have been used as evidence for a domain-general structure processor is that those manipulations activated a domain-general component of Broca’s area. This component belongs to the domain-general multiple demand (MD) network implicated in executive control and goal-directed behaviors and robustly sensitive to effort (e.g., Duncan, 2010, 2013; Fedorenko et al., 2013). Manipulations of hierarchical complexity have often been confounded with difficulty, such that the more structurally complex conditions required greater cognitive effort. As a result, they would be likely to elicit responses in the MD network, including its inferior frontal component residing in Broca’s area (see Fedorenko & Blank, in press, for more discussion). Although it is possible that outside of the domain of language— which appears to rely on domain-specific processing mechanisms (Fedorenko et al., 2011)—this component of the MD network, or the MD network as a whole, is important for structured behavior, we should keep in mind that this network is also sensitive to manipulations that don’t involve structured/hierarchical representations (e.g., Crittenden & Duncan, 2012). The latter findings argue against the idea of complex syntactic operations being the core computation of the MD network.

***Third***, we have here focused on language comprehension. Could the architecture of language processing be different for language *production*? Although we plausibly access the same knowledge representations to interpret (comprehend) and generate (produce) linguistic utterances—in line with substantial overlap that has been observed between comprehension and production in fMRI (e.g., Menenti et al., 2011; Silbert et al., 2014)— the computational demands of language production differ from those of language comprehension. In particular, the goal of comprehension is to infer the intended meaning from the linguistic signal, and abundant evidence now suggests that the representations we extract and maintain during comprehension are probabilistic and noisy (e.g., Ferreira et al., 2002; Levy et al., 2008; Gibson et al., 2013). In contrast, in production, the target meaning is (typically) clear and precise, and the goal is to express that particular meaning. To do so, we have to utter a precise sequence of words where each word takes a particular morpho-syntactic form, and the words appear in a particular order. This pressure for linearization of words, morphemes, and sounds might lead to a clearer temporal, and perhaps spatial, segregation among the different stages of the production process compared to comprehension (e.g., Hagoort & Indefrey, 2014), and/or require additional mechanisms implemented in regions that do not support comprehension. Indeed, recent evidence from intracranial stimulation suggests that a small region in posterior superior temporal cortex may be selective for encoding and enacting morpho-syntactic inflections (Lee, Fedorenko, et al., 2018; see Fedorenko et al., 2018, for discussion). It is therefore possible that some aspects of language production are implemented in focal and functionally selective regions. However, this conjecture remains to be evaluated further in future work.

***Finally***, in the current study, we investigated the relationship between syntactic and lexico-semantic processing in a single Germanic language: English. Some of the theoretical linguistic work, experimental psycholinguistic work, and computational modeling work have spanned multiple languages (e.g., Norcliffe et al., 2015). However, much/most cognitive neuroscience research has been conducted on English and a handful of other languages/families (e.g., see Bornkessel-Schlesewsky & Schlesewsky, 2016 for discussion). Thus, the conclusions drawn here remain to be generalized to typologically diverse languages.

### Other findings that have been interpreted as evidence for syntax selectivity

Three other lines of research—on phenomena that are, or have been, taken as strong evidence for syntax selectivity—deserve discussion. ***First***, the early ERP literature on language processing appeared to have provided evidence of distinct components associated with lexico-semantic processing (N400; Kutas & Hilliyard, 1980) vs. with syntactic processing (P600; Osterhout & Holcomb, 1992; Hagoort et al., 1993). However, the interpretation of the P600 as an index of syntactic processing has been challenged from the earliest days following its discovery (e.g., Coulson et al., 1998), and the current dominant interpretation of this component is as a domain-general error detection or correction signal (e.g., Kolk & Chwila, 2007; Vissers et al., 2007; van de Meerendonk et al., 2010; Sassenhagen et al., 2014; Ryskin et al., 2020b). Some other, earlier, ERP components (e.g., eLAN; Friederici, 2002) have been argued to index syntactic processes. However, the robustness and nature of these components have been questioned (Steinhauer & Drury, 2012), and the interpretation that seems to capture the empirical findings best has to do with violations of word-form expectations rather than syntactic parsing per se (e.g., Dikker et al., 2009, 2010; Rosenfelt et al., 2009, 2011).

***Second***, syntactic priming—re-use of a syntactic frame based on recent linguistic experiences (Bock, 1986; see Pickering and Ferreira, 2008; Branigan & Pickering, 2016 for reviews)—has often been cited as evidence of abstract syntactic representations independent of meaning (e.g., Bock & Loebell, 1990), including in relatively recent cognitive neuroscience papers (Pallier et al., 2011). However, a large body of work has now established that the effect is strongly modulated by lexical overlap (e.g., Mahowald et al. 2016; Scheepers et al., 2017; Ziegler et al., 2019) and driven by the meaning-related aspects of the utterance (e.g., Hare & Goldberg, 1999; Cai et al., 2012; Ziegler & Snedeker, 2018; Ziegler et al., 2018).

And ***third***, a class of linguistic phenomena known as “syntactic islands” (Ross, 1967) have been argued to be due to abstract properties of syntactic structures unrelated to meaning. In particular, some structures are disfavored when a phrase is “extracted” from its “canonical” structural location in a sentence (e.g., Chomsky, 1973; Schütze et al., 2015). However, this interpretation has been challenged: in particular, some researchers (e.g., Erteschik-Shir, 1973; Kuno, 1987; Goldberg, 2013) have argued that semantic and discourse factors can explain these phenomena (see Abeillé et al., in press, for empirical support).

### Beyond syntax and semantics: dissociations of other linguistic processes

Language processing encompasses a broad array of computations in both comprehension and production, and some aspects of language are robustly dissociable and supported by distinct sets of brain regions. Here, we have argued that during language comprehension the mechanisms that process the *structure* of sentences are also deeply sensitive to the meanings of *individual words*. Is the reverse also true? Are there any brain areas that process individual word meanings but are not sensitive to syntactic processing? One area that deserves a mention lies in the left temporal pole, extending onto the lateral and ventral surface of the temporal lobe. According to one influential hypothesis, this area has been implicated in lexical retrieval (e.g., Damasio et al., 2004; Drane et al., 2008; Grabowski et al., 2001; Tranel, 2006, 2009; Mesulam et al., 2013, 2015). According to another hypothesis, motivated chiefly by investigations of semantic dementia (Gorno-Tempini et al., 2011), this region has been linked to general object knowlege (e.g., Lambon-Ralph et al., 2001; Rogers et al., 2004, 2006; Patterson et al., 2007). Critically, syntactic abilities in patients with damage to this area appear to be relatively preserved (e.g., Mesulam et al., 2013, 2015). This area overlaps with our LAntTemp fROI, but extends beyond it. In our experiments, the LAntTemp language fROI showed stronger responses during lexico-semantic than syntactic processing in two of the three experiments (**Figure 5**). However, this region still responds reliably to syntactic processing in at least some manipulations: for example, it responds more strongly to a) structured but meaningless Jabberwocky sentences compared to lists of unconnected nonwords (Fedorenko et al., 2010; see Mollica et al., in prep., for a replication), and b) structurally more complex sentences with object-extracted relative clauses compared to those with subject-extracted relative clauses (Blank et al., 2016). Further, based on evidence from MEG, parts of the left anterior temporal lobe have been implicated in semantic composition, above and beyond the processing of single words (e.g., Bemis & Pylkkanen, 2011; see Pylkkanen, 2019 for a review). The brain imaging evidence therefore suggests that this region is engaged in some syntax-/combinatorics-relevant processes, in addition to lexico-semantic processing. But because the temporal pole and the anterior ventral temporal cortex are challenging to study with fMRI given the signal dropout due to proximity to air-filled sinuses (e.g., Devlin et al., 2000), it is possible that we are missing some areas—anterior to our LAntTemp fROI and/or on the ventral surface of the temporal lobe—that are truly selective for lexico-semantic/conceptual over syntactic processing. The patient evidence mentioned above implies the existence of such areas, although the evidence is not unequivocal (e.g., Bi et al., 2010). More work is therefore needed to functionally characterize left anterior temporal areas and their relationship with the core fronto-temporal language network.

Going beyond syntax and semantics, lower-level speech perception and reading processes as well as speech production (articulation) recruit areas that are robustly distinct from the high-level areas that we focused on here. In particular, speech perception recruits parts of the auditory cortex in the superior temporal gyrus and sulcus (e.g., Scott et al., 2000, Mesgarani et al., 2014; Overath et al., 2015), and these areas are highly selective for speech over many other types of auditory stimuli (Norman-Haignere et al., 2015). Reading recruits a small area on the ventral surface of the temporal lobe (see McCandliss et al., 2003, for a review), and this “visual word-form area” is highly selective for letters in a familiar script over a broad range of other visual stimuli (Baker et al., 2007; Hamame et al., 2013). And articulation draws on a set of areas, including portions of the precentral gyrus, supplementary motor area, inferior frontal cortex, superior temporal cortex, and cerebellum (e.g., Wise et al., 1999; Bohland and Guenther, 2006; Eickhoff et al., 2009; Basilakos et al., 2017).

On the other end of linguistic processes, discourse-level processing draws on areas distinct from those that support word and sentence-level comprehension (e.g., Ferstl & von Cramon, 2001; Lerner et al., 2011; Jacoby et al., 2018; Blank & Fedorenko, 2020), and aspects of non-literal language have been argued to draw on brain regions in the right hemisphere (e.g., Joanette et al., 1990) and on the system that supports social cognition (e.g., Kline et al., 2018; Hagoort, 2019).

Thus, many aspects of language are robustly dissociable, in line with distinct patterns of deficits reported in the aphasia literature (e.g., Goodglass, 1993). However, syntactic and lexico-semantic processing do not appear to be separable during language comprehension. Some brain areas not easily accessible to fMRI may be selective for lexico-semantic processing, as discussed above, but no area within the language network appears to be selective for syntactic processing based on both brain imaging studies and patient investigations.

## Concluding remarks

To conclude, across three fMRI experiments, we found robust responses to both lexico-semantic and syntactic processing throughout the language network, with generally stronger responses to lexico-semantic processing, and no regions, or even sets of non-contiguous voxels within those regions, that respond reliably more strongly to syntactic processing than lexico-semantic processing. These results constrain the space of possible neural architectures of language. In particular, they rule out architectures that postulate a distinct region (or set of regions) that selectively supports syntactic/combinatorial processing (i.e., architectures shown in **Figures 1a-d**). These findings, illuminating how minds are instantiated in brains, are mirrored by studies of how minds are implemented in machines, where modern-day connectionist networks achieve remarkable performance on a wide variety of language tasks (e.g., Mikolov et al., 2010; Sutskever et al., 2014; Bahdanau et al., 2016), including those that involve complex syntactic phenomena (e.g., Linzen et al., 2016; Gulordava et al., 2018; Futrell et al., 2018, 2019; Prasad et al., 2019; Wilcox et al., 2019a,b), apparently without a clearly separable syntax-selective mechanism. Taking all the available data into consideration, it therefore seems that a cognitive architecture whereby *syntactic processing is not separable from the processing of individual word meanings* is most likely.

## Author contributions

EF conceived and designed the study. All authors collected the data and performed first-level data analyses. IB performed second-level analyses. IB, EF, and MS created the figures. EF drafted the manuscript and IB provided critical revisions, with MS and ZM providing additional comments. All authors approved the final version of the manuscript.

## Acknowledgements

We would like to acknowledge the Athinoula A. Martinos Imaging Center at the McGovern Institute for Brain Research at MIT, and its support team (Steve Shannon and Atsushi Takahashi). We thank former and current EvLab members (especially Zuzanna Balewski and Brianna Pritchett) for their help with data collection, Gina Kuperberg for providing the materials used in adapted form in Experiment 1, Michael Behr for creating the script for Experiment 1, Zuzanna Balewski for help with creating the materials and script for Experiment 2, Josef Affourtit for helping put together the OSF page, Nancy Kanwisher for discussions of the experimental design for all three experiments, and Inbal Arnon, Yonatan Belinkov, Ted Gibson, Adele Goldberg, Maryellen MacDonald, and Jayden Ziegler for comments on the manuscript. We also thank the audience at the 2017 CUNY Sentence Processing conference (Cambridge, MA) for feedback, as well as three anonymous reviewers whose comments helped to greatly improve the manuscript. EF was supported by NIH awards R00-HD057522, R01-DC016607, R01-DC016950, by a grant from the Simons Foundation to the Simons Center for the Social Brain at MIT, and by support from the Brain and Cognitive Sciences Department and the McGovern Institute for Brain Research at MIT.

## Conflict of interest

The authors declare no competing financial interests.

## Footnotes

1 One type of manipulation missing here is one that relies on ambiguity. Both lexical (e.g., Rodd et al., 2005, 2010, 2012; Davis et al., 2007; Mason and Just, 2007; Zempleni et al., 2007; Bekinschtein et al., 2011) and structural (e.g., Mason et al., 2003) ambiguity have been investigated in prior fMRI studies—although the former has received more attention—and both have been shown to elicit responses in the inferior frontal and posterior temporal cortex. Because, to the best of our knowledge, no arguments for syntactic selectivity have been made based on stronger responses to syntactic than lexical ambiguity in some part(s) of the language network, we did not include ambiguity manipulations here.

2 It is worth noting that, similar to the Lexico-semantic and Syntactic conditions, the Global meaning condition also elicited a response that was reliably stronger than the Same condition in all language fROIs (*p*s<0.05). This effect provides evidence that language regions are sensitive to differences in complex meanings above and beyond the meanings of individual words (given that the only thing that differs between the sentences in a pair in the Global-meaning condition is word order).

